# Multiomic single-cell lineage tracing to dissect fate-specific gene regulatory programs

**DOI:** 10.1101/2022.10.23.512790

**Authors:** Kunal Jindal, Mohd Tayyab Adil, Naoto Yamaguchi, Xue Yang, Helen C. Wang, Kenji Kamimoto, Guillermo C. Rivera-Gonzalez, Samantha A. Morris

## Abstract

Complex gene regulatory mechanisms underlie differentiation and reprogramming. Contemporary single-cell lineage tracing (scLT) methods use expressed, heritable DNA barcodes to combine cell lineage readout with single-cell transcriptomics enabling high-resolution analysis of cell states while preserving lineage relationships. However, reliance on transcriptional profiling limits their adaptation to an ever-expanding tool kit of multiomic single-cell assays. With CellTag-multi, we present a novel approach for independently profiling lineage barcodes with single-cell chromatin accessibility without relying on co-assay of transcriptional state, paving the way for truly multiomic lineage tracing. We validate CellTag-multi in mouse hematopoiesis, characterizing transcriptional and epigenomic lineage priming across progenitor cell populations. In direct reprogramming of fibroblasts to endoderm progenitors, we use CellTag-multi to comprehensively link early cell state with reprogramming outcomes, identifying core regulatory programs underlying on-target and off-target reprogramming. Further, we reveal the Transcription Factor (TF) Zfp281 as a novel regulator of reprogramming outcome, biasing cells towards an off-target mesenchymal fate via its regulation of TGF-β signaling. Together, these results establish CellTag-multi as a novel lineage tracing method compatible with multiple single-cell modalities and demonstrate its utility in revealing fate-specifying gene regulatory changes across diverse paradigms of differentiation and reprogramming.

## Introduction

The quantification of cell identity is crucial to understanding development, disease, and homeostasis, yet the notion of cell identity remains poorly defined^1^. Single-cell technologies, now tailored to diverse modalities^2^, are expanding our understanding of how cell identity is established and maintained^3^. In particular, single-cell lineage tracing (scLT) methods allow cell relationships to be tracked throughout biological processes, revealing cell fate decisions during differentiation and reprogramming^4, 5^. Prospective scLT methods label cells with unique genetic ‘barcodes’ that are expressed as RNA; capturing these barcodes via single-cell RNA-seq (scRNA-seq) allows the parallel capture of lineage information and single-cell transcriptomes^6–13^.

These methods to barcode and track cells have been deployed across several *in vitro* differentiation and reprogramming paradigms^5, 14^. The accessibility of cells within these systems permits longitudinal sampling and cellular barcoding at precise time points, allowing early progenitor state to be linked to terminal fate (termed ‘state-fate analysis’; **Fig. 1a**). Such a strategy has been used to determine how well gene expression state in progenitors reflects eventual cell fate in hematopoiesis^13^. This work demonstrated that subsequent fate could be predicted, albeit with limited accuracy, from progenitor gene expression, indicating the existence of heritable fate determinants that are not captured by scRNA-seq alone. Similarly, viral barcoding, ‘CellTagging,’ of transcription factor-mediated direct reprogramming of mouse embryonic fibroblasts (MEFs) to induced endoderm progenitors (iEPs), suggested that reprogramming outcome is determined during the early stages of fate conversion^7^. However, the early gene regulatory changes that set cells on their destined path have not been fully characterized. Additional information from epigenomic assays such as single-cell Assay of Transposase Accessible Chromatin by sequencing (scATAC-seq) may be crucial to uncover the heritable properties that play a key role in the establishment and maintenance of cell identity. Previously, natural DNA variation has been used to infer coarse cellular phylogenies with scATAC-seq^15, 16^. However, the resolution of such retrospective methods is limited due to their reliance on the accrual of somatic mutations. In contrast, the density of lineage information recorded can be precisely controlled at biologically relevant time points using successive rounds of cellular barcoding^7, 17^ with prospective methods. This is essential for profiling early, lineage-specific responses in dynamic systems like differentiation and reprogramming.

**Figure 1.**
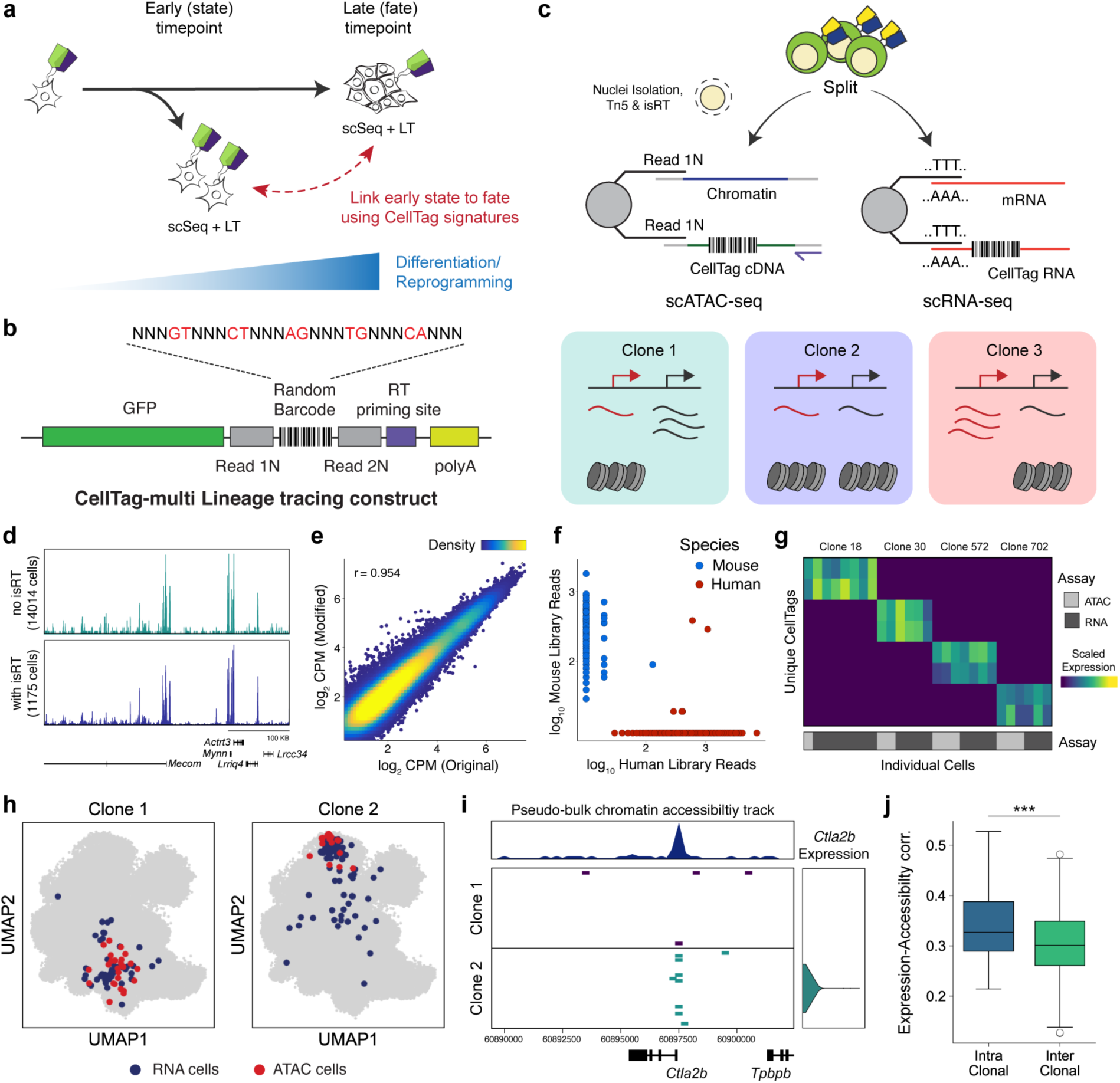
CellTag-multi allows simultaneous capture of lineage information with gene expression and chromatin accessibility. (**a**) A framework for relating early cell state with fate using single-cell lineage tracing. (**b**) Schematic depicting the CellTag-multi lineage tracing construct. (**c**) Schematic detailing parallel capture of CellTags during scRNA-seq and modified scATAC-seq library preparation, using targeted isRT of CellTags in intact nuclei. CellTag-multi enables simultaneous clonal tracking of transcriptional and epigenomic state. (**d**) Browser tracks comparing chromatin accessibility signal across aggregated scATAC-seq profiles generated using the original and modified library preparation methods. (**e**) Scatterplot comparing log normalized reads in ATAC peaks across aggregated scATAC-seq profiles generated with the original and modified library preparation methods. (r = Pearson Correlation Coefficient). (**f**) Plot for the human-mouse species mixing experiment depicting the number of CellTag reads/cell from each CellTag library. (**g**) Heatmap showing scaled CellTag expression in scRNA-seq and scATAC-seq sisters for four multiomic clones identified in a population of expanded reprogramming fibroblasts. (**h**) Joint UMAP of RNA and ATAC cells with two clones (clone 1 and clones 2) cells projected, along with assay information. (**i**) Browser track showing single-cell accessibility at the *Ctla2b* locus and *Ctla2b* gene expression across clones 1 and 2. Top Panel: Pseudo-bulk accessibility signal at the *Ctla2b* locus. (**j**) Box plots comparing intra- and inter-clonal correlation between clonally aggregated gene expression and gene activity scores in the reprogramming dataset (n = 62 clones used; Mann Whitney Wilcoxon test; p-value = 5.39e-4).

To enable prospective lineage tracing with chromatin accessibility capture, we have developed ‘CellTag-multi.’ CellTag-multi is based on our previous CellTagging technology, which uses sequential lentiviral delivery of CellTags (heritable random barcodes) to enable the construction of multi-level lineage trees^7, 17^. Here, we introduce a new strategy in which CellTags, expressed as polyadenylated transcripts, can be captured in both scRNA-seq and scATAC-seq assays allowing for independent tracking of clonal transcriptional and epigenomic state.

We validate this method using *in vitro* hematopoiesis, a well-characterized model of multi-lineage differentiation, and demonstrate highly accurate reconstruction of lineage relationships and capture of lineage-specific progenitor cell states across scRNA-seq and scATAC-seq. Moreover, the addition of chromatin accessibility information to gene expression allows for a significant improvement in the prediction of differentiation outcome from early progenitor state. We also deploy CellTag-multi in the direct lineage reprogramming of fibroblasts to induced endoderm progenitors (iEPs), to characterize early genomic events in rare subpopulations of cells that successfully reprogram. This application reveals how chromatin is remodeled following expression of reprogramming TFs, enabling deeper insight into gene regulatory network reconfiguration. We uncover the TF Foxd2 as a facilitator of on-target reprogramming, increasing the efficiency of MEF to iEP conversion. Conversely, we identify Zfp281 as a TF biasing cells towards an off-target mesenchymal fate via its regulation of TGF-β signaling, which we validate experimentally. We demonstrate that the identification of these TFs as novel reprogramming regulators is only possible via multiomic profiling. Together, these findings highlight the utility of CellTag-multi in defining the molecular regulation of early cell state and its relation to fate across diverse biological applications.

## Development and validation of CellTag-multi

CellTagging relies on single-cell capture of CellTags — heritable DNA barcodes expressed as polyadenylated transcripts^7, 17, 18^. In the standard workflow, CellTags are captured as transcripts and reverse transcribed (RT), along with cellular mRNA, during 3’ end scRNA-seq library preparation. In contrast, scATAC-seq directly captures fragments of the accessible genome, omitting capture of CellTag transcripts, rendering CellTagging incompatible with scATAC-seq assays. To enable CellTag profiling with scATAC-seq, we introduced two essential modifications. First, we developed an *in situ* Reverse Transcription (isRT) step to selectively reverse transcribe CellTag barcodes inside intact nuclei. By introducing this additional step after transposition, we omitted the need to RT CellTags during scATAC-seq library construction. Second, we modified the CellTag construct to flank the random barcode with Nextera Read 1 and Read 2 adapters (**Fig. 1b, Ext Fig. 1a, b**).

During scATAC-seq library preparation, nuclei are partitioned into nanoliter droplets along with single-cell barcoding beads and PCR reagents. Each bead contains a barcoded forward primer complementary to the Nextera Read 1 adapter to barcode and linearly amplify all ATAC fragments during the GEM incubation step. By inserting Nextera Read 1 and Read 2 adapters in the CellTag construct, we enabled single-cell capture of reverse transcribed CellTags along with accessible chromatin during the GEM incubation stage (**Fig. 1c, Ext Fig. 1b**). This strategy improved the CellTag capture rate by >200-fold compared to the unmodified scATAC-seq protocol (**Ext Fig. 1c**). Additionally, we introduced a reverse primer specific to the CellTag cDNA during GEM incubation to exponentially amplify CellTag fragments, while ATAC fragments undergo linear amplification (**Supplementary Table 1, Ext Fig. 1b**). Together, these modifications led to a >50,000-fold increase in CellTag capture (**Ext Fig. 1c**), with CellTags being detected in >96% of cells in scATAC-seq relative to 98% in scRNA-seq (**Ext Fig. 1d**), without negatively impacting scATAC-seq data quality or genome-wide chromatin accessibility signal (**Fig. 1d, e, Ext Fig. 1e, f**).

To support the accurate identification of clonally related cells, it is essential that CellTag signatures from individual cells are captured with high fidelity, minimizing background noise. To assess the fidelity of CellTag signatures captured in scATAC-seq, we performed a species-mixing experiment (**Ext Fig. 2a**). We labeled human (HEK 293T) cells and mouse (expanded iEPs) cells with two different versions of the CellTag-multi library, combined nuclei isolated from both populations in a 1:1 ratio and profiled them using our modified scATAC-seq method. Plotting CellTag reads/cell, we observed that nuclei from each species predominantly consisted of reads from the expected CellTag library, indicating minimal inter-species crosstalk (**Fig. 1f; Ext Fig. 2b, c**).

Finally, to perform large-scale lineage tracing experiments, we synthesized a complex CellTag-multi library containing ∼80,000 unique barcodes, as confirmed by sequencing (**Methods**). We applied CellTag-multi to a population of expanded mouse fibroblasts undergoing reprogramming to iEPs and profiled clones with scRNA-seq and scATAC-seq, detecting CellTags in 70% (RNA) and 51% (ATAC) of the cells at an average MOI of 2 (RNA) and 2.5 (ATAC). Filtering, error-correction, and allowlisting of CellTag reads (**Methods**) enabled high-fidelity identification of distinct clones across the two single-cell modalities (**Fig. 1g, h, Ext Fig. 2d-f**). As expected, the correlation between gene expression and accessibility was higher within clones than across clones (**Fig. 1i, j**). These analyses established the efficacy of CellTag-multi for the labeling and capture of clonally related cells across scRNA and scATAC modalities. Next, we leveraged CellTag-multi to link early state with cell fate in diverse cell fate specification and reprogramming paradigms.

## Benchmarking CellTag-multi using an *in vitro* model of mouse hematopoiesis

To validate lineage analysis across single-cell modalities with CellTag-multi, we applied it to hematopoiesis, a well-characterized paradigm for multi-lineage differentiation. Recently, scLT was used to define the early transcriptional cell states that lead to defined differentiation outcomes in mouse hematopoiesis. However, these analyses suggested that early transcriptional changes alone cannot fully define future cell fate and posited a role for cell states that evade transcriptional profiling, collectively termed hidden state variables^13^. In this context, we aimed to apply CellTag-multi to further refine state-fate linkages in early hematopoiesis by identifying fate-specific changes in both early gene expression and chromatin accessibility.

We isolated Lin^-^, Sca1^+^, c-Kit^+^ (LSK) cells from adult mouse bone marrow and cultured them in broad myeloid differentiation media^13^. Upon isolation, we tagged these cells with the CellTag-multi library to track clones across modalities. To capture both early state and fate across clones, we profiled half of the cells 60 hours after initiation of differentiation (Day 2.5; state sample), re-plated the remaining cells across two technical replicates, and collected them for sequencing on Day 5 (fate sample). In the case of both samples, cells were split between scRNA-seq and scATAC-seq (**Fig. 2a**), resulting in the profiling of 9,789 state cells (scRNA-seq: n=5,161; scATAC-seq: n=4,628) and 67,029 fate cells (scRNA-seq: n=56,534; scATAC-seq: n=10,495 cells), after quality filtering (**Ext Fig. 3a, b**). We identified cells from all major hematopoietic lineages across single-cell modalities (**Fig. 2b, Ext Fig. 3c**). CellTagging was consistent across single-cell modalities, yielding 83-99% labeled cells.

**Figure 2.**
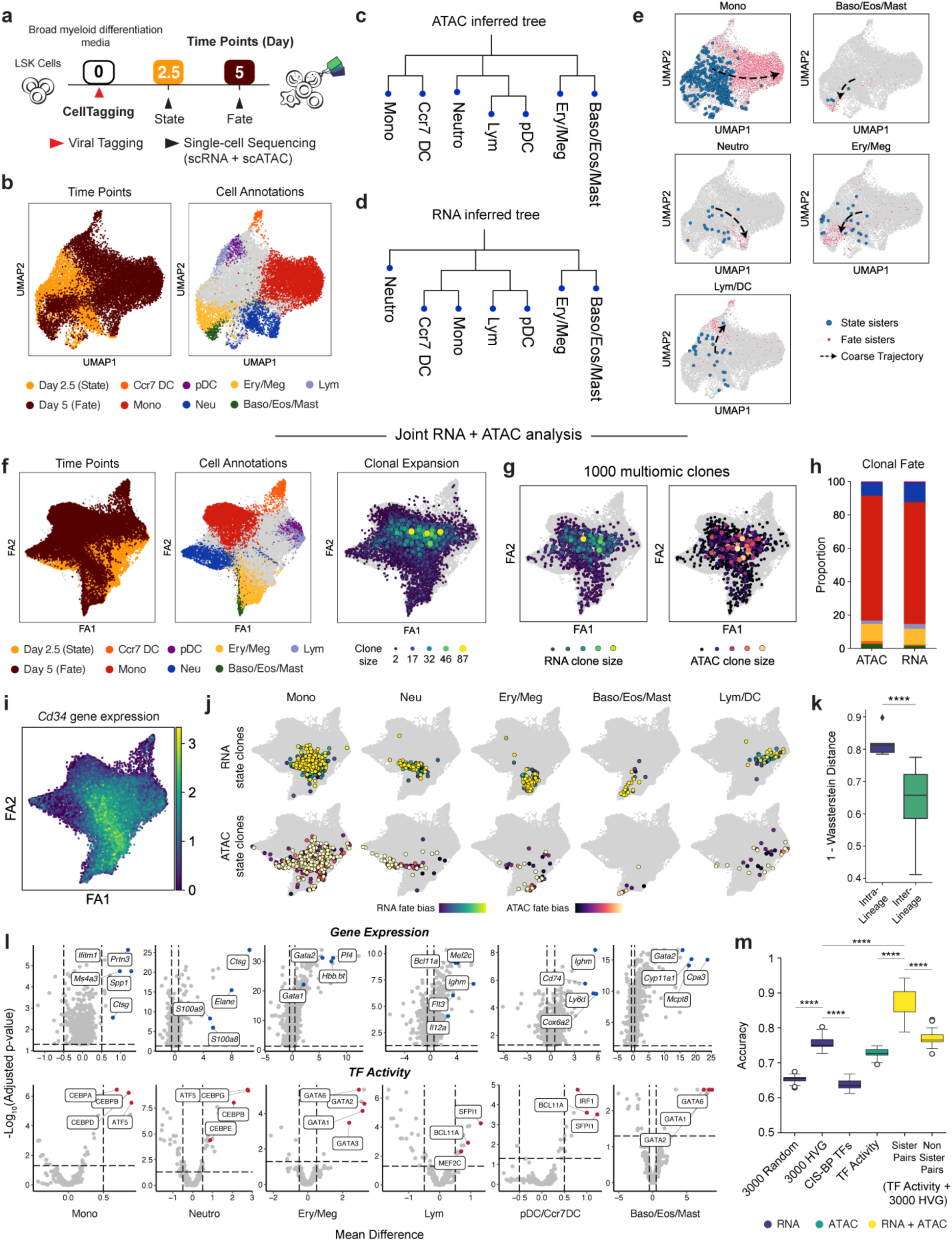
Application of CellTag-multi to link early hematopoietic cell state with fate. (**a**) Schematic detailing the experimental design for the *in vitro* hematopoiesis state-fate experiment. (**b**) scATAC-seq UMAPs with time point (left panel) and cell fate information (right panel) projected (Mono: Monocytes; Neu: Neutrophils; Lym: Lymphoids; Ery: Erythroids; Meg: Megakaryocytes; Baso: Basophils; Eos: Eosinophils; Mast: Mast Cells; pDC: plastoid Dendritic Cells). Only major cell fates are highlighted. Hematopoietic lineage hierarchy as inferred from (**c**) scATAC or (**d**) scRNA clone coupling. (**e**) scATAC-seq UMAPs with state and fate sisters for major hematopoietic fates highlighted. (**f**) Clone-cell embedding UMAPs with time point and cell fate information projected onto cells (left and center panels) and clonal expansion information projected onto clones (right panel), detailed cell type annotations are shown in **Ext Fig. 4c.** (**g**) UMAPs with RNA and ATAC clonal expansion information projected onto a thousand random multiomic clones. Both modalities display biased expansion of early myeloid cells, consistent with our differentiation culture conditions. (**h**) Bar plot depicting distribution of cell fates across RNA and ATAC clones (Fates are colored as in Fig. 2b). (**i**) UMAP with scaled *Cd34* expression level, a marker of Hematopoietic Stem and Progenitor Cells (HSPCs), projected onto the scRNA cells. (**j**) UMAPs with state (Day 2.5) sub-clones for each major lineage highlighted along the differentiation continuum for both single-cell modalities, with fate bias information projected. (**k**) Box plot comparing overlap between RNA and ATAC state sub-clones within and across cell fates (Mann Whitney Wilcoxon test; p-value = 3.76e-5). (**l**) Volcano plots summarizing the results of differential feature enrichment analysis for each group of state sub-clones across for scRNA (top panel) and scATAC modalities (bottom panel). (**m**) Box plot summarizing accuracy scores of trained state-fate prediction models. Machine learning partially predicts cell fate from Day 2.5 state across both modalities. However, predictive performance increases significantly when both are considered together, highlighting the existence of unique functional priming in both gene expression and chromatin accessibility state (Mann Whitney Wilcoxon test; p-values: **** = p < 0.0001, HVG: Highly Variable Genes, n = 25 accuracy values for each model (**Methods**)).

To compare clonal analysis across modalities, we first analyzed the scRNA-seq and scATAC-seq datasets separately and identified clones in each modality independently (**Ext Fig. 3d**). Lineage hierarchies inferred using clonally related cells (**Methods**) were consistent across scRNA and scATAC despite the chromatin dataset comprising fewer cells, demonstrating the ability of CellTag-multi in defining fate relationships using clonal scATAC-seq data alone (**Fig. 2c, d**). Assigning a fate label to each clone, based on the most abundant cell type amongst its Day 5 sisters, allowed mapping of coarse fate trajectories on the 2D embeddings (**Fig. 2e, Ext Fig. 3e**). Joint clone calling across both datasets led to an increase in number of cells tracked (**Ext Fig. 3f**), likely due to clones that are split across modalities (multiomic clones). We identified a total of 37,441 scRNA-seq cells in 5,973 clones and 6,098 scATAC-seq cells in 3,012 clones, labeled with 4.2 CellTags/cell (in scRNA-seq) and 3.4 CellTags/cell (in scATAC-seq) on average (**Ext Fig. 3g, h**). 2,227 clones spanned both state and fate samples, including 877 multiomic clones. These clones were used for the remainder of the analyses.

For visualization, we co-embedded cells from both modalities using Canonical Correlation Analysis (CCA)^19^. Further, we devised a unique clone-cell co-embedding approach to include clones as individual data points in a single-cell embedding, enabling straightforward visualization and assessment of clone-level metadata and global trends across clones (**Ext Fig. 3i)**. We first extracted the cell-cell similarity graph, produced as part of standard single-cell analysis workflows. In this graph, each cell is represented by a node and the connection between a pair of cells is weighted based on their phenotypic similarity. Next, we imputed abstract clone nodes and clone-cell edges to this graph based on clonal data. Finally, we used this expanded clone-cell graph as input for dimensionality reduction algorithms such as UMAP^20^ or ForceAtlas^21^ to produce a single 2D-embedding of the data, where both cells and clones are represented by individual points. We applied this visualization to the hematopoiesis data to co-embed RNA and ATAC cells with all clones, with minimal impact on the underlying structure of the data (**Fig. 2f, Ext. Fig. 3j, k**). Clones, now represented as individual data points, faithfully represented their constituent cells (**Ext Fig. 3l)** and can be used to visualize clonal metadata across all cells (**Fig. 2f, right panel**). Consistent with previous reports, we observe continuous transitions from progenitor populations to distinct hematopoietic lineages across modalities, as previously reported^13, 22, 23^ (**Ext Fig. 4a-c**). While CellTag capture was uniform across cell states (**Ext Fig. 4d**), we observed higher clonal expansion along the monocyte lineage, consistent with our myeloid differentiation culture conditions (**Fig. 2f right panel, g**).

**Figure 3.**
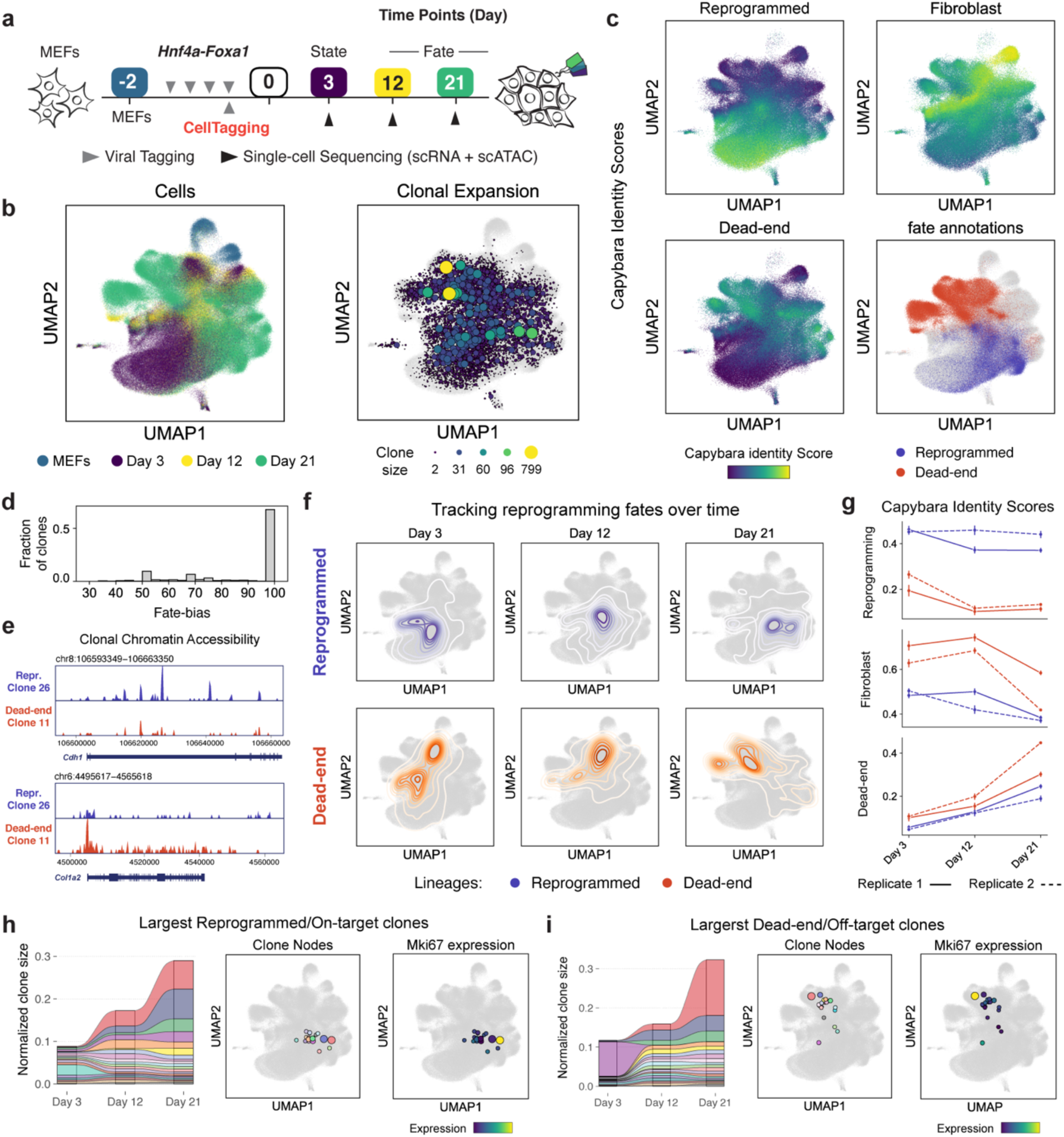
Application of CellTag-multi to dissect clonal fate dynamics in direct reprogramming. (**a**) Experimental design for the direct reprogramming state-fate experiment. (**b**) Cells from both scRNA-seq and scATAC-seq, across all time points, were co-embedded with clones and visualized using a UMAP. (Left Panel) Time point information projected on cells. (Right Panel) Clonal expansion visualized using clone nodes. (**c**) Capybara transcriptional identity scores projected on scRNA-seq cells for reprogrammed, dead-end and fibroblast cell identities, based on a previous lineage tracing dataset^7^. Cell fates were annotated for Days 12 and 21. Reprogrammed and dead-end cell fates are highlighted (Lower Right Panel). (**d**) Histogram of fate bias across all state-fate clones. (**e**) Clonal chromatin accessibility browser tracks for two dead-end and reprogramming clones. (**f**) Contour plots showing longitudinal tracking of cell fates enabled by CellTagging. (**g**) Transcriptional identity dynamics tracked along both lineages. Dead-end cells depart from a MEF like identity and acquire an off-target reprogrammed state. Significant clonal expansion is observed along both lineages, as depicted via alluvial plots, clone nodes and clonal expression levels of *Mki67* (a proliferation marker gene) in the 20 largest (**h**) reprogramming/on-target and (**i**) dead-end/off-target clones.

We linked Day 2.5 cell state with Day 5 fate, by re-assigning each clone, from the joint clone calling results, a fate label based on the most abundant cell type amongst its Day 5 sisters (**Fig. 2h**, **Ext Fig. 4e**). To map early clonal state along the differentiation continuum, we extended our clone-cell embedding approach further and split each clone into sub-clones (up to four) based on the assay and time point capture of each sister (**Ext Fig. 4f**). While Day 5 fate sub-clones localized largely within their respective cell fate clusters (**Ext Fig. 4g**), Day 2.5 state sub-clones associated with each major fate formed distinct groups closer to the undifferentiated progenitors (**Fig. 2i, j**), suggesting early functional priming of immature cells. Moreover, state sub-clones within the same ‘fate potential’ group overlapped significantly across single-cell modalities (Mann Whitney Wilcoxon test; p-value = 3.76e-5, **Fig. 2j, k**), demonstrating high-fidelity capture of state-fate linkages across transcriptional and epigenomic states with CellTag-multi. Projecting fate bias scores, defined as the fraction of fate sisters belonging to the assigned clonal fate, on to state sub-clones, we observed that low fate bias clones occupied areas closer to the overlapping boundaries of each fate potential region, likely indicating areas of multi-potency (**Fig. 2j, Ext Fig. 4h**).

To characterize these fate-specific changes in early cell state on a molecular level, we assessed the enrichment of transcriptional and epigenetic signatures in Day 2.5 sisters for each fate group (**Fig. 2l; Methods**). Using gene expression, we identified several known fate-specific markers in each group, such as *Spp1*^13^ and *Ms4a3*^24^ in the Monocyte primed group; *Elane* and *Ctsg*^13^ in the Neutrophil primed group; *Pf4*^25^ and *Gata2*^13^ in the Erythroid/Megakaryocyte groups. In the Lymphoid group, we identified *Flt3*, a predominantly Lympho-myeloid gene^26^, and several lymphoid fate-specific genes such as *Mef2c*^27^ and *Bcl11a*^28^. For epigenetic data, we focused on TF activity scores^29^, which estimate the enrichment of TF motifs in single-cell epigenomes^29^. Unlike peak accessibility, TF activity feature space is dense and continuous, allowing comparison between small groups of cells, and is easier to interpret relative to individual peak features^29^. TF activity enrichment analysis revealed several expected lineage specifying TFs for each fate^22, 30^, such as several CEBP TFs enriched in Monocyte and Neutrophil primed groups; GATA1 and GATA2 in the Erythroid/Megakaryocyte and Basophils/Eosinophils/Mast cells groups; Lympho-myeloid TF SFPI1 (also known as PU.1) in the Lymphoid and Dendritic Cells (DC) group, along with BCL factors and MEF2 factors, indicating extensive epigenomic priming in early cells towards their respective cell fate. A complete list of differential gene expression and TF activity enrichment can be found in Supplementary Table 2.

## Chromatin accessibility and gene expression jointly define fate predictive cell state

Our above state-fate analysis suggests that lineage-specific changes in gene expression are accompanied by extensive epigenetic remodeling, rendering the genome more accessible to fate-specifying TFs. Previous analysis has suggested that cell states hidden from transcriptional profiling play a role in fully defining fate-associated changes in cell state^13^. Changes in chromatin accessibility could account for some of this hidden variance and we tested this hypothesis by assessing whether cell fate can be accurately predicted from early state using our multiomic clonal data.

We trained machine learning models to predict clonal cell fate from gene expression or chromatin accessibility profiles of Day 2.5 sisters (**Ext Fig. 5a**). We tested three different architectures: Logistic Regression, Random Forest, and LightGBM, and assessed model performance using prediction accuracy (**Ext Fig 5b**). Overall, Random Forest models performed the best and were used for all downstream analysis. For gene expression, we trained a classification model to predict clonal fate using expression of the three thousand most highly variable genes (HVG) and obtained an accuracy of 75.6% (**Fig. 2m, Ext Fig. 5c**). For chromatin accessibility, we used Day 2.5 imputed TF activity scores (**Methods**) for 884 TF motifs to predict clonal fate and obtained an accuracy of 72.7% (**Fig. 2m**). Notably, an RNA model trained on expression levels of TFs, obtained from the Catalog of Inferred Sequence Binding Preferences (CIS-BP) database, only scored only 63.8% on prediction accuracy (**Fig. 2m**). The significantly lower predictive performance of TF expression compared to TF activity could be attributed to either technical dropout in scRNA-seq or significantly higher lineage specific priming of TF binding sites compared to TF expression, or a combination of both.

To assess fate-specific priming in different functional regions of the epigenome, we computed TF activity scores using subsets of accessible peaks and compared fate prediction performance across these feature spaces. Specifically, we computed TF activity scores using only promoter, distal, exonic, or intronic peaks and trained fate prediction models with each. We observed significant variation in performance between different ATAC models, indicating different levels of fate-specific epigenetic priming across functional regions of the genome (**Ext Fig. 5d**). This variation was independent of the number of peaks used to compute each set of TF activity scores (**Ext Fig. 5d**). Distal and Intronic were the best performing models, comparable in performance to the full peak set model (‘All’). Promoter and Exonic models performed significantly worse, suggesting that fate-specifying epigenetic changes during these early stages were dominated by changes in distal regulatory regions of the epigenome rather than accessibility of genes themselves. This observation is reinforced by the persistence of TF enrichment trends across state groups in distal and intronic subsets but not in the exonic and promoter subsets (**Ext Fig. 5e**). We confirmed these results using SHAP, a game theoretic approach to quantify the contributions of individual input features in explaining the output of a machine learning model^31^. Indeed, SHAP analysis showed that in the better-performing models, an increase in CEBP/A motif accessibility and an increase in MECOM motif accessibility were better predictors of Monocyte and Ery/Meg fates, respectively, suggesting a lack of functional priming in the promoter-proximal accessible genome (**Ext Fig. 5f, g**).

Finally, we tested whether combining RNA and ATAC features is more predictive of fate than either individual modality. For this, we trained a combined RNA and ATAC model where RNA and ATAC Day 2.5 sister cells within the same clone were paired randomly, and their combined gene expression and TF activity signatures were used to predict clonal fate label. This analysis was limited to multiomic state-fate clones. The combination of both state modalities was significantly better at predicting fate (mean accuracy score = 86.5%) compared to either individual modality or pairs of unrelated RNA and ATAC state cells (**Fig. 2m**). These results show that both gene expression and chromatin accessibility jointly comprise cell states that define future cell fate. Moreover, these modalities consist of non-redundant and highly complementary state information, as a combination of both predicts cell fate much more accurately than each modality in isolation.

## Dissecting clonal dynamics of direct reprogramming

Our application of CellTag-multi to hematopoiesis demonstrated the method’s utility to capture informative gene regulatory dynamics in a well-characterized differentiation paradigm. We next applied CellTag-multi to a less defined system — the direct reprogramming of MEFs to iEPs driven by retroviral overexpression of Hnf4*α* and Foxa1^7, 32, 33^. Direct lineage reprogramming presents a unique paradigm of cell identity conversion, with cells often transitioning through progenitor-like states or acquiring off-target identities^34, 35^. Such non-linear fate dynamics can be challenging to assess, especially when relying solely on the computational inference of cell fate trajectories^12^. Ground truth lineage tracing serves as a crucial resource for dissecting lineage-specific cell state changes during direct reprogramming^7^. Originally reported to yield hepatocyte-like cells^32^, we have previously shown that Hnf4*α* and Foxa1 overexpression in MEFs generates cells with the broader potential to functionally engraft liver and intestine^18, 33, 36^. This prompted their re-designation as ‘induced Endoderm Progenitors’ (iEPs). More recently, we have further characterized the similarity of long-term cultured iEPs to regenerating Biliary Epithelial Cells (BECs)^37^.

Using our original CellTag-based lineage tracing, we identified two distinct iEP reprogramming trajectories: a successful ‘reprogrammed’ trajectory, characterized by endodermal and hepatic gene expression, and a ‘dead-end’ trajectory, defined by a failure to extinguish the starting fibroblast identity^7^. Further work demonstrated key functional differences between these fates, with successfully reprogrammed cells harboring the potential to engraft acutely damaged mouse intestine^18^. Our previous lineage tracing suggests that the reprogrammed and dead-end fates are determined in the early stages of fate conversion^7^. However, our original CellTagging methodology did not capture any epigenetic information and only sparsely sampled early state clones, limiting mechanistic insight into these initial reprogramming stages.

Here, we deployed CellTag-multi in iEP reprogramming, modifying our clonal resampling strategy to optimize state-fate analysis (**Fig. 3a**). First, we transduced MEFs with Hnf4*α* and Foxa1 for 48 hours to initiate reprogramming, in two independent biological replicates. During the last 12 hours of this 48-hour period, we transduced cells with the complex CellTag-multi library, enabling clonal relationships to be tracked. 72 hours following the final viral transduction (Reprogramming Day 3), we collected two-thirds of the cells for single-cell RNA and ATAC profiling (state sample) and re-plated the remaining cells. Subsequent samples were collected on Days 12 and 21 (fate samples) to assess reprogramming outcome. We also profiled the starting MEF population (scATAC-seq, this study; scRNA-seq from a previous study^7^)), resulting in a total of 466,459 single-cells (scATAC-seq: 223,686; scRNA-seq: 242,863) in the final dataset after quality filtering (**Ext Fig. 6a, b**). We identified a total of 8,502 clones, containing 46,438 cells (Replicate 1: 3,068 clones; Replicate 2: 5,416 clones, average clone sizes of 4.8 and 5.9 cells/clone, respectively (**Ext Fig. 6c, d)**). We identified 1,428 ‘state-fate’ clones across both replicates, defined as clones that spanned state (Day 3) and at least one fate time point, Day 12 or Day 21 (**Ext Fig. 6d**).

Following dimensionality reduction and clustering of the co-embedded RNA and ATAC datasets, clone-cell embedding was performed (**Fig. 3b, Ext Fig. 6e, f, g**). We annotated Day 12 and 21 fate clusters (‘reprogrammed,’ ‘dead-end,’ and ‘transition’) based on expression and accessibility of known reprogramming associated genes, and unsupervised cell-type classification based on transcriptional state using Capybara^37^ (**Fig. 3c; Ext Fig. 7a, b**). In line with our previous reports^7, 18, 37, 38^, reprogrammed cells express epithelial and iEP markers, *Cdh1* and *Apoa1*, respectively. Dead-end cells are characterized by the retention of fibroblast gene expression but are still transcriptionally distinct from MEFs, expressing low levels of iEP markers and several dead-end-specific genes such as *Sfrp1,* a Wnt signaling modulator^7^ (**Ext Fig. 7b, c**). Transition cells represent states in between MEFs and reprogrammed/dead-end identities. Following cluster annotation, we assigned fate labels to each state-fate clone. As the majority of state-fate clones showed high fate-bias, we assigned clonal fate based on the most abundant cell annotation amongst the fate sisters (**Fig. 3d**), identifying 1,009 reprogrammed, 2,493 dead-end and 1,371 transition clones. Dead-end and reprogrammed clones displayed a lineage-specific increase in accessibility of known marker genes (**Fig. 3e**).

Using clonal information, we linked each reprogrammed and dead-end clone to its Day 3 state sisters, allowing us to track changes in cell identity longitudinally (**Fig. 3f**). These results were consistent when clonal analysis was performed for each modality independently (**Ext Fig. 7d-f**). Comparing Capybara transcriptional cell identity scores across lineages, we found that iEP identity scores were consistently higher along the reprogrammed lineage compared to the dead-end lineage. MEF identity scores, while significantly higher along the dead-end lineage, exhibited a steep decline after Day 12 coinciding with an increase in dead-end transcriptional identity score (**Fig. 3g**). This suggested a delayed departure from MEF identity to an alternate cell state. We observed high levels of clonal expansion along both lineages (**Fig. 3h, i**). These observations suggest that despite retaining expression of canonical fibroblast marker genes, dead-end cells are a fundamentally distinct, off-target cell state and reprogramming outcome. Thus, the ‘reprogrammed’ and ‘dead-end’ fates are better described as ‘on-target’ and ‘off-target’ reprogramming, respectively.

## Linking early state to fate reveals molecular features of off-target reprogramming

Next, to identify early state changes that regulate entry onto distinct fate trajectories, we focused on Day 3 state clones destined to on-target (reprogrammed) or off-target (dead-end) reprogramming fates. From assessing the distribution of Day 3 sisters destined to either of the two fates, it is evident that they are not localized to defined clusters (**Ext Fig. 8a, b**). Further, trajectory inference using CellRank^39^ fails to reveal these initial states (**Ext Fig. 8c**), demonstrating the importance of ground truth lineage tracing. We found that both Day 3 gene expression and TF activities were highly predictive of clonal fate. Similar to our analysis of hematopoiesis, fate prediction accuracy was significantly higher when both modalities were considered, as compared to either modality individually. Further, distal and intronic peaks were more predictive of fate than proximal and exonic (**Ext Fig. 8d, e**).

To identify early molecular signatures of lineage specification, we compared gene expression, chromatin accessibility, and TF activity scores across MEFs and Day 3 state sisters grouped by fate outcome. Comparing gene expression enrichment across the three groups, 2,116 genes were differentially enriched with 1,582 enriched genes uniquely defining each group (**Fig. 4a, Ext Fig. 8f**). While some genes displayed transient fate-specific expression, others consistently increased over time in a lineage-specific manner (**Supplementary Table 3**). Early iEP marker genes such as *Apoa1* were enriched in both on- and off-target trajectories on Day 3, consistent with our previous observation that most cells initiate reprogramming^7^ (**Ext Fig. 8f, g**). On-target (reprogrammed) enriched genes included *Krt19*, a marker of BECs, Wnt signaling associated genes *Wnt4*, *Anxa8,* and epithelial marker *Ezr* (**Fig. 4b, Supplementary Table 4**). Top off-target (dead-end) related genes included canonical smooth muscle markers *Acta2* and *Tagln* and other mesenchymal genes such as *Ptn,* and *Ncam1*, suggesting broad engagement of mesenchymal programs, in addition to *Sfrp1*, a Wnt signaling pathway inhibitor (**Fig. 4b, Supplementary Table 4**).

**Figure 4.**
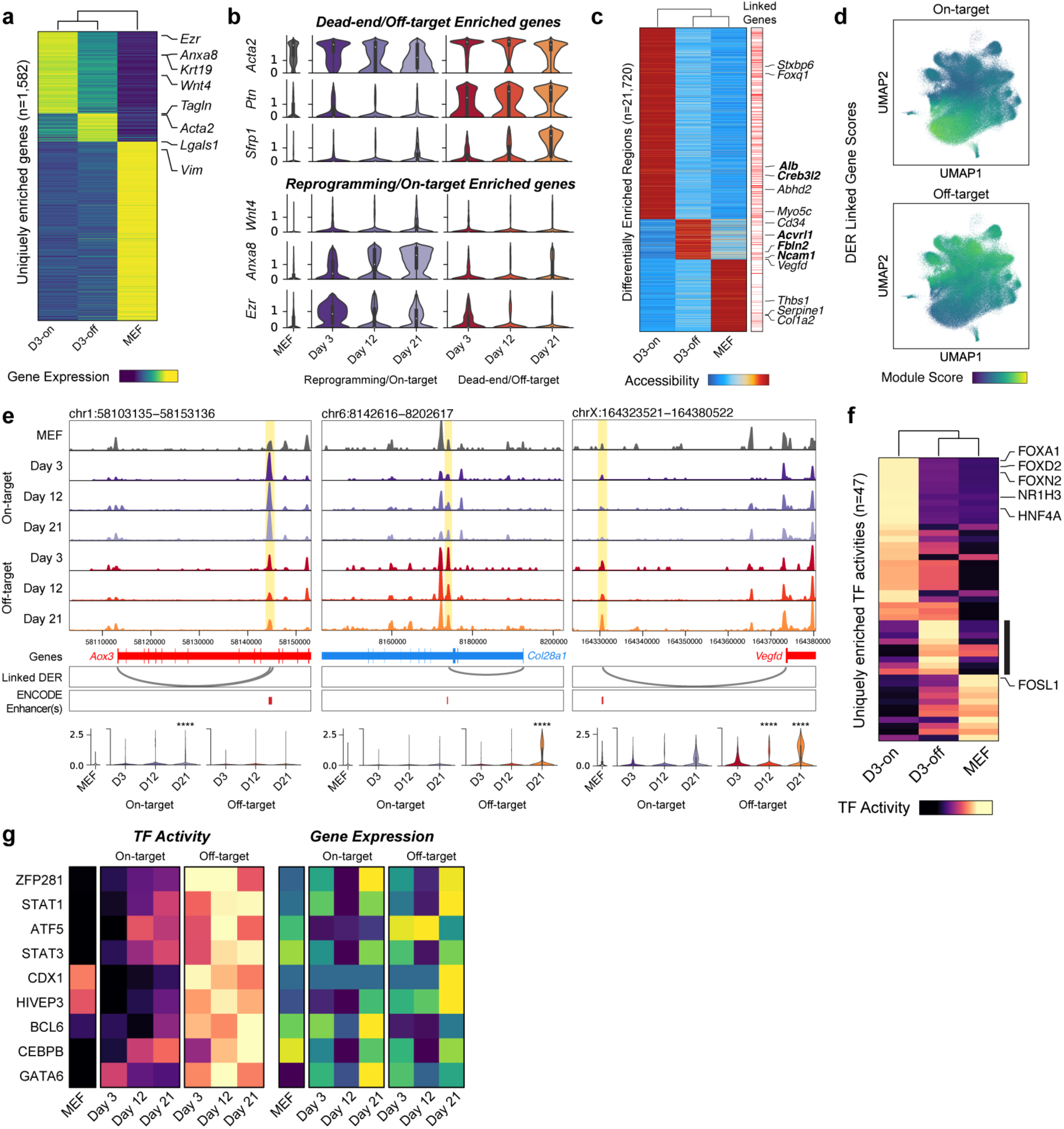
Assessing fate-specific changes in early cell state. (**a**) Heatmap of genes uniquely enriched across uninduced MEFs or one of the two reprogramming fates on Day 3 (FDR threshold: 0.05, log fold change threshold: 0). (**b**) Violin plots of several genes enriched in either off-target (dead-end) destined or on-target (reprogramming) destined cells. (**c**) Heatmap of peaks uniquely enriched across uninduced MEFs or one of the two reprogramming fates on Day 3 (FDR threshold: 0.05, log fold change threshold: 1). Right panel shows annotation of peaks linked to genes (Methods). (**d**) Module scores for genes linked to either on-target or off-target DERs projected onto the clone-cell embedding. (**e**) Top panel: Accessibility browser tracks for each lineage split by day, highlighting peaks linked to late lineage markers (On-target: *Aox3*; Off-target: *Col28a1* and *Vegfd*) showing lineage specific changes in accessibility on Day 3. The *Aox3* and *Vegfd* linked DERs overlap perfectly with an ENCODE enhancer like element (ELS) while the *Col28a1* linked DER is within 100 bp of an ELS. Bottom panel: Expression levels of the three genes across MEFs and the two reprogramming lineages split by days. The asterisks (*) mark time points and lineage of significant differential enrichment. (**f**) Heatmap of TF activities uniquely enriched across uninduced MEFs or one of the two reprogramming fates on Day 3 (FDR threshold: 0.05, mean difference threshold: 0.5). (**g**) Left Panel: Heatmap showing TF activity (left panel) and gene expression (right panel) levels for off-target associated TFs in MEFs and each reprogramming lineage split by time points. TF activity signatures show a much stronger lineage bias as compared to gene expression values.

Comparing genome-wide chromatin accessibility revealed 21,720 Differentially Enriched Regions (DERs) across Day 3 on-target and off-target destined cells and uninduced MEFs, indicating extensive fate-specific epigenetic reconfiguration during early reprogramming (**Fig. 4c, Supplementary Table 5**). DERs were enriched for distal and intergenic peaks, suggesting epigenetic re-patterning of distal regions as a driver of cell fate conversion, consistent with our above observations in hematopoiesis (**Ext Fig. 8h**). Motif analysis revealed enrichment of reprogramming and hepatic TFs in on-target DERs, and several TFs with documented roles in mesenchymal fates^40, 41^ in off-target DERs (**Ext Fig. 8i, j**). Using our paired RNA and ATAC data, we linked accessible peaks to genes and identified 37,058 putative cis-regulatory elements (CREs)^42^ (**Fig 4c**, **Methods**). Gene-linked peaks were enriched for enhancer-like signatures (ELS) from the ENCODE candidate CRE database^43^ (**Methods**, **Ext Fig. 8k**). Genes linked to on-target and off-target DERs displayed fate-specific expression patterns (**Fig. 4d, Ext Fig. 8l)**. On-target DERs consisted of several CREs linked to endodermal genes, such as *Alb*, *Foxq1,* and *Creb3l2.* In contrast, off-target DERs contained CREs linked to mesenchymal genes such as *Ncam1*, a modulator of Mesenchymal Stromal Cell migration^44^, *Fbln2*, a mesenchymal gene associated with embryonic heart development^45^, and *Vegfd*, a regulator of angiogenesis^46^ and endothelial differentiation of bone marrow-derived mesenchymal stem cells^47^ (**Fig. 4c; Supplementary Table 5**). In several instances, this analysis captured lineage-specific changes in accessibility of CREs before significant changes in gene expression were detected. For instance, a *Vegfd*-linked CRE overlapping with an ENCODE enhancer displayed enrichment in dead-end destined cells (Day 3), while expression changes were not detectable until Day 12. Similar regulatory changes were observed for *Aox3*^48^, a liver-associated aldehyde oxidase, and *Col28a1,* an oligodendrocyte enriched collagen^49^, prior to changes in gene expression (**Fig. 4e, Supplementary Table 5**).

To identify functional changes in chromatin accessibility on a genomic scale, we compared inferred TF activities across on-target and off-target destined cells and uninduced MEFs. To preclude potential false positives, we discarded all TFs with low correlation (< 0.3) with their respective gene activity scores, identifying 47 uniquely enriched TFs (**Fig. 4f, Ext Fig. 8m, Supplementary Table 6**). On-target destined cells were highly enriched for the two reprogramming TFs, FOXA1, and HNF4A. Other on-target associated TFs included FOXD2, FOXO1, and NR1H3, a hepatic fate-specifying TF^50^ (**Fig. 4f**). We identified a set of nine TFs uniquely enriched in off-target destined cells (**Fig. 4f (black bar), g**). Several of these TFs (Zfp281, Cebpb, Gata6, Hivep3) have been previously documented to play a role in regulating mesenchymal cell identities^51–54^. Surveying the expression data, none of the off-target TFs display a similar fate-biased enrichment (**Fig. 4g, Ext Fig. 8n**), highlighting the importance of lineage-specific chromatin profiling in identifying these targets. This lack of enrichment could be due to technical dropout during scRNA-seq or due to secondary mechanisms regulating the genomic engagement of these TFs.

Altogether, our lineage-specific multiomic assessment of iEP generation demonstrates clear early molecular differences associated with reprogramming outcomes. Indeed, from as early as reprogramming day 3, cells on the dead-end lineage exhibit unique characteristics. Rather than retaining MEF identity, we observe that the dead-end lineage constitutes a highly proliferative, mesenchymal cell state with unique markers and regulatory changes, thus representing an ‘off-target’ reprogrammed state. The early specification of this state is supported by our GRN inference using CellOracle^38^, suggesting that network reconfiguration is unique to each trajectory and is established early in the reprogramming process. CellTag-multi has the potential to define the molecular features of these early states, offering deeper mechanistic insight into the reprogramming process.

## Foxd2 and Zfp281 as drivers of on- and off-target reprogramming

Higher accessibility of both motifs and genomic targets^55^ of FOXA1 and HNF4A in on-target cells on Day 3 suggests significant differences in genomic engagement of the reprogramming TFs between the two fate outcomes (**Fig. 5a, Ext Fig. 9a**). This could, at least in part, be explained by differential expression levels of the *Hnf4α-Foxa1* transgene across the two lineages, with off-target destined cells displaying significantly lower transgene expression (**Fig. 5a**; Mann Whitney Wilcoxon test; p-value = 6.5e-42). However, we have also previously described an off-target trajectory expressing high transgene levels, suggesting additional mechanisms influencing genomic engagement by the reprogramming TFs^38^.

**Figure 5.**
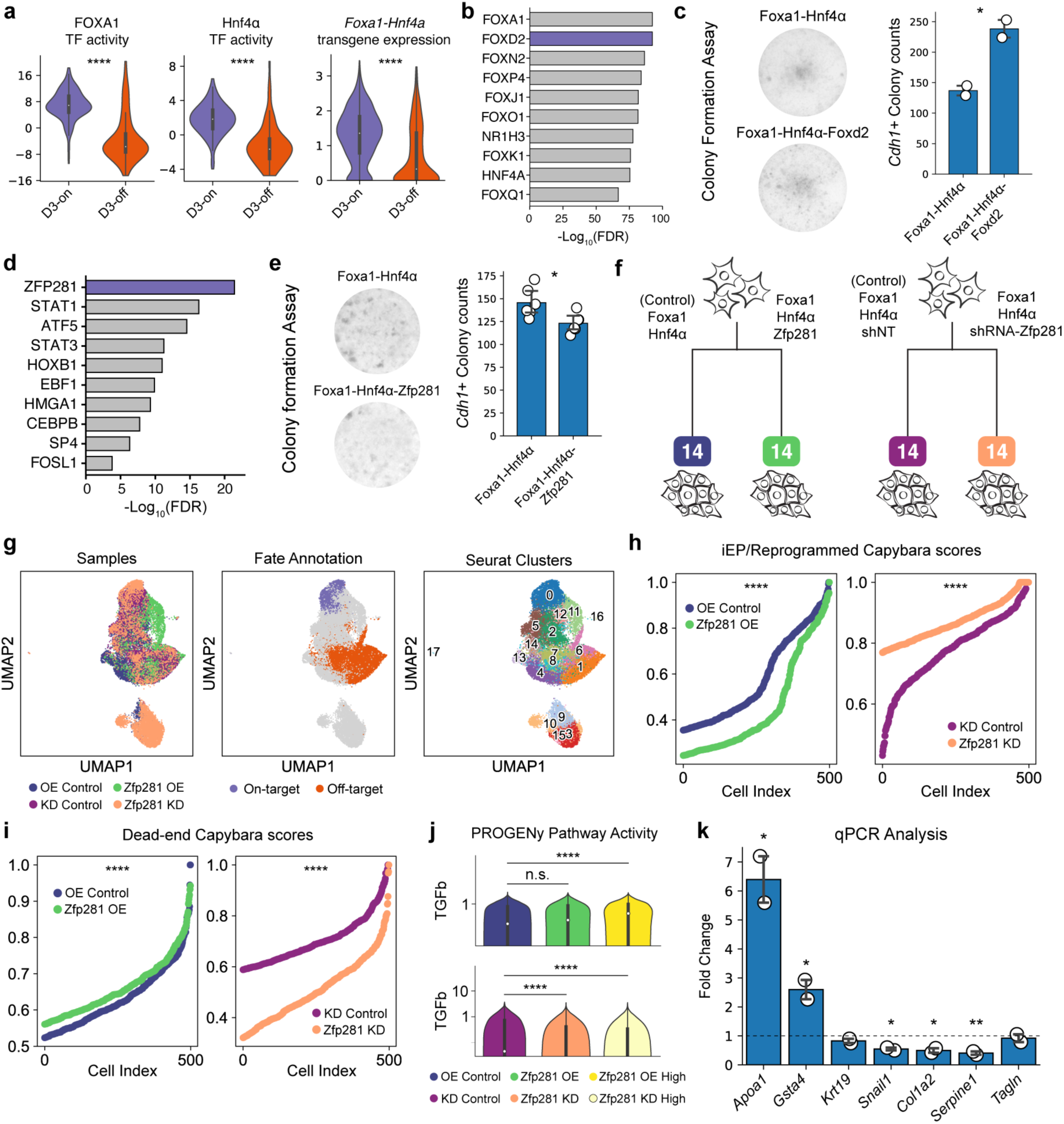
Identification of TF regulators of on-target and off-target reprogramming fate. (**a**) Left and Middle panels: Violin plots comparing enrichment of FOXA1 and HNF4A TF activities across the two reprogramming fates on Day 3 (Mann Whitney Wilcoxon test; p-values: FOXA1 = 9.2e-22, HNF4A = 1.7e-20). Right panel: Violin plot comparing enrichment of the *Hnf4α-Foxa1* transgene expression across the two reprogramming fates on Day 3 (Mann Whitney Wilcoxon test; p-value = 6.5e-42). (**b**) Top ten TFs enriched in on-target destined cells based on TF activity scores. (**c**) Left Panel: Representative images from the *Foxd2* overexpression colony formation assay; Right Panel: Bar plot showing increase in CDH1-positive colony counts in *Foxd2* overexpressing cells compared to a standard reprogramming experiment (t-test; p-value = 0.045; n = 2 biological replicates). (**d**) Top ten TFs enriched in off-target destined cells based on TF activity scores. (**e**) Left Panel: Representative images from the *Zfp281* overexpression colony formation assay; Right Panel: Bar plot showing decrease in CDH1-positive colony counts in the *Zfp281* overexpressing sample compared to a standard reprogramming experiment. (t-test; p-value = 0.017; n = 6 biological replicates). (**f**) Schematic of the scRNA-seq experiment for *Zfp281* over-expression (OE) and knockdown (KD) during reprogramming. A GFP expression vector and non-target shRNA were used as controls for OE and KD respectively. (**g**) UMAP embedding for all cells profiled in the *Zfp281* OE and KD experiments with sample information (Left), cell fate annotation (Middle) and Seurat cluster information (Right) projected. (**h**) Plot of iEP Capybara identity scores across the KD and OE samples compared to respective controls (Mann Whitney Wilcoxon test; p-values: *Zfp281* OE vs control = 1.07e-53; *Zfp281* KD vs control = 2.19e-53). (**i**) Plot of dead-end Capybara identity scores across the KD and OE samples compared to respective controls (Mann Whitney Wilcoxon test; p-values: *Zfp281* OE vs control = 1.11e-11; *Zfp281* KD vs control = 3.26e-120). (**j**) Violin plots showing variation of TGF-β pathway activity across control vs OE vs OE high cells (upper panel) and control vs KD vs KD high cells (lower panel). OE high cells are defined as the subset OE sample cells with above average *Zfp281* expression. KD high cells are defined as the subset of KD sample cells with below average *Zfp281* expression. (**k**) Bar plots showing fold-change in reprogramming and dead-end marker genes upon small molecule mediated inhibition of TGF-β signaling, compared to a vehicle control, on Day 5 of iEP reprogramming (t-test; p-values: *Apoa1* = 0.02, *Col1a2* = 0.02, *Gsta4* = 0.04, *Serpine1* = 0.009, *Snail1* = 0.01; n=2 technical replicates).

Outside of FOXA1, and HNF4A, we identified FOXD2 as the top on-target fate-specifying TF candidate (**Fig. 5b, Ext Fig. 9b**). Adding Foxd2 to the Foxa1 and Hnf4*α*reprogramming cocktail led to significantly increased expression of the iEP marker *Cdh1* and decreased expression of mesenchymal marker *Tagln* on reprogramming day 12 (t-test; p-values: *Cdh1* = 0.03; *Tagln* = 0.006; 2 biological replicates; 2 technical replicates **Ext Fig. 9c**). In addition, colony formation assays showed a significant increase in the number of CDH1-positive colonies formed with the addition of Foxd2 to the standard iEP reprogramming cocktail (t-test; p-value=0.045; 2 biological replicates; **Fig. 5c**), validating its role in improving on-target fate conversion.

The top off-target-enriched candidate was ZFP281, a Zinc Finger protein (**Fig. 5d, Ext Fig. 9d**). Zfp281 is a known regulator of cell fate in mouse embryonic stem cells^56^ and promotes epithelial-to-mesenchymal transitions^57^. To further confirm the inferred enrichment of ZFP281 TF activity in off-target fated cells, we performed Tomtom motif similarity analysis^58^ to identify TFs that share a motif similar to ZFP281. We found four other TF motifs that were both significantly similar to the ZFP281 motif (adjusted p-value < 0.05) and were enriched in off-target destined cells. Amongst these TFs, ZFP281 displayed the highest enrichment in the off-target lineage both in terms of gene expression and TF activity (**Ext Fig. 9e**). Additionally, single-cell accessibility of ZFP281 genomic targets^56^ was positively correlated with inferred ZFP281 TF activity (Pearson’s correlation coefficient = 0.53; **Ext Fig. 9f**) and ZFP281 regulated genes^59^ were significantly more predictive of cell fate as compared to a size-matched set of random genes (Mann Whitney Wilcoxon test; p-value = 2.248e-09; **Ext Fig. 9g**), further confirming its role in off-target fate specification during iEP reprogramming. Notably, both *Zfp281* and *Foxd2* failed to show a strong lineage-specific bias in gene expression levels, highlighting the unique insights offered by multiomic lineage tracing in the identification of fate-specifying TFs (**Ext Fig. 9h**).

Indeed, inclusion of Zfp281 along with Foxa1 and Hnf4*α* in the reprogramming cocktail resulted in a moderate but statistically significant reduction in the number of CDH1-positive colonies (t-test; p-value = 0.017; **Fig. 5e**). To further characterize the role of Zfp281 in reprogramming, we performed both overexpression (OE) and shRNA mediated knockdown (KD) of Zfp281, along with respective control samples, and profiled cells with single-cell sequencing on reprogramming day 14 (**Fig. 5f, g, Ext Fig. 10a**). We found that the rate of reprogramming (both on- and off-target) increased with increasing *Zfp281* expression (**Ext Fig 10b**), suggesting a role for Zfp281 in accelerating fate conversion in iEP reprogramming. Moreover, we identified a distinct subpopulation of cells, predominantly consisting of Zfp281 KD cells that were depleted for expression of key markers of both on-target and off-target reprogramming such as *Apoa1* and *Ctla2a* (**Ext Fig 10c-e**). These cells were enriched for MEF and early off-target marker gene expression, depleted for both off-target and on-target markers genes from Day 21 (obtained from our lineage analysis; **Ext Fig. 10f-h**) and thus likely represent a “stalled” cell state due to reduced *Zfp281* expression levels. Despite its acceleration of cell fate conversion broadly, we found that Zfp281 shifted the identity of reprogrammed cells away from an iEP-like state and towards a dead-end/off-target-like state consistently across the OE and KD experiments (**Fig. 5h, i**), confirming a role for Zfp281 in biasing cells towards an off-target fate, as suggested by our lineage tracing analysis. This finding also explains the reduced number of CDH1-positive colonies observed in our colony formation assay, despite the increase in total number of on-target reprogrammed cells upon Zfp281 overexpression.

Finally, a key downstream effector of Zfp281 is TGF-β signaling^51, 60^, an Epithelial-to-Mesenchymal Transition (EMT) associated pathway^61^. Indeed, TGF-β pathway activity, as inferred using PROGENy^62^ (**Methods**), increased with *Zfp281* OE and decreased with *Zfp281* KD, suggesting active regulation of TGF-β signaling by Zfp281. Given that on-target reprogramming is characterized by cellular epithelialization and off-target reprogramming is characterized by activation of broad mesenchymal programs, we hypothesized that increased TGF-β signaling mediated via Zfp281 acts as a barrier to on-target reprogramming. Indeed, inhibition of TGF-β signaling during iEP reprogramming using the small molecule SB431542^63^ led to a significant increase in expression of reprogramming marker genes *Apoa1* and *Gsta4* and a significant decrease in mesenchymal/off-target genes such as *Serpine1, Snail1, Col1a2* (**Fig. 5k**). This was accompanied by an increase in epithelial/iEP-like morphology as early as day 3 of reprogramming (**Ext Fig 10i**) suggesting a crucial role for TGF-β signaling, downstream of Zfp281, in determining fate outcome during iEP reprogramming.

## Discussion

Here we have presented CellTag-multi, a method for independent single-cell lineage tracing across scRNA-seq and scATAC-seq assays. In the context of hematopoiesis, we have used CellTag-multi to map transcriptional and epigenomic states of progenitor cells and link them to clonal fate, recapitulating enrichment of known lineage-specific cell state signatures across progenitor populations. With chromatin state, we showed that lineage-specific epigenetic priming is associated with changes in accessibility of known fate-specifying TF motifs and that such changes occur primarily in the regions of the genome distal to promoters. Previous analysis has demonstrated the inability of early transcriptional state alone in predicting cell fate and posited a role for alternate cell state modalities^13^. By exploiting multiomic clonal relationships, we demonstrated that the predictability of cell fate from state is significantly improved when both early transcriptional and epigenomic state are considered, as opposed to either modality individually, suggesting that the RNA and ATAC modalities consist of non-redundant and highly complementary state information.

Our application of CellTag-multi to the less characterized paradigm of iEP reprogramming generated similar observations, where multiomic clonal data captured in the early stages of fate conversion is highly predictive of reprogramming outcome. Again, fate-specifying epigenetic changes during early stages of differentiation are dominated by changes in distal regulatory regions of the epigenome. Further, we have been able to molecularly characterize the ‘dead-end’ state as a highly proliferative, mesenchymal-like cell state, representing an ‘off-target’ reprogrammed state. Indeed, a similar state has been reported in direct reprogramming of mesenchymal stromal cells to induced hepatocytes, revealing the appearance of *Acta2*- expressing mesenchymal cells during the reprogramming process^64^. Outside of the hepatic lineage, off-target identities have been reported in other reprogramming paradigms^35, 65^, suggesting that this may be a more general feature of lineage reprogramming.

Our multiomic lineage tracing demonstrates the establishment of on- and off-target trajectories from early stages, supported by our earlier transcriptome-based lineage tracing of iEP reprogramming^7^ and GRN inference^38^. However, given the single modality capture of relatively few clones in that earlier study, we were not able to comprehensively characterize early molecular states. Here, the collection of ground truth data on lineage, transcriptome, and epigenome has allowed us to better characterize these distinctive early states, enabling novel mechanistic insights into reprogramming. We have shown crucial early differences in gene regulation that lead to distinct reprogramming outcomes. Specifically, we have identified and experimentally validated that Foxd2 promotes successful reprogramming, while Zfp281 activity leads to engagement with an off-target trajectory. Differences in reprogramming TF levels may account for these early differences. However, lower levels of exogenous TF expression do not simply lead to reprogramming failure, as the off-target fate is molecularly unique from fibroblasts and could be considered a reprogramming byproduct in itself. These results suggest that the stoichiometry of TF overexpression in these reprogramming models may offer further insight into how TFs control cell identity. Single-cell analysis of TF binding could provide further insights into the role of differential binding of the two reprogramming TFs in specifying off-target fate.

Our recovery of Foxd2 and Zfp281 as novel regulators of early-stage reprogramming was not possible from differential gene expression analysis alone, demonstrating the utility of CellTag-multi. Our data suggests off-target enriched Zfp281 activity from early stages of reprogramming. From our experimental validation, we found that knockdown of Zfp281 expands a population of cells in a ‘stalled’ state, where they fail to extinguish fibroblast gene expression while upregulating off-target cells. Conversely, overexpression of Zfp281 helps accelerate fate conversion, resulting in a considerable increase in reprogramming efficiency. However, Zfp281 still draws the reprogrammed cells toward an off-target, mesenchymal-like state. A role for this TF in driving broad mesenchymal expression programs, including components of the TGF-β signaling pathway, has recently been described^51^. Here, we demonstrate that the inhibition of TGF-β signaling enhances on-target marker expression while decreasing off-target gene expression. These results suggest a potential strategy to enhance on-target reprogramming, where Zfp281 expression can help erase the starting cell identity while blocking downstream TGF-β signaling might prohibit entry onto the off-target trajectory.

Altogether, the data we present here across two distinct biological systems demonstrates that lineage-specific capture of gene expression and chromatin accessibility provides rich information on gene regulation, offering unique mechanistic insights into the specification and maintenance of cell identity. More widely, single-cell lineage tracing has revealed distinct, clonally heritable transcriptional states across various biological systems^66–68^. These phenotypic differences, arising from seemingly non-genetic sources, have strong biological implications. For example, clonal variability in cell state has been shown to impact malignant clonal expansion and efficacy of drug treatment in cancer cells^66, 68^. Elsewhere, CRISPR-based systems have been used to create mutable barcodes to allow multi-level lineage recording without the need for successive rounds of cell labelling^69, 70^. Given its versatility and ease of use, we envision that CellTag-multi can be readily applied to such biological questions and use cases.

Finally, we have developed CellTag-multi to work independently with scRNA-seq and scATAC-seq, as existing single-cell methods that co-assay multiple modalities from the same cell^71–74^ can suffer from lower data quality compared to methods that profile each modality individually. Further, enabling the capture of lineage in parallel with chromatin accessibility provides users with additional flexibility for experimental design. Advances in single-cell technologies are allowing measurement of an ever-increasing number of cellular modalities. A similar expansion in lineage tracing assays will complement these new methods with multiomic, clonal tracking of cell state and enable deeper mechanistic insight into the regulation of cell identity and clonal heritability of cell state. CellTag-multi, with its cell lineage read out alongside gene expression and chromatin accessibility, paves the way for such multiomic, single-cell lineage tracing methods.

## Acknowledgments

We thank members of the Morris laboratory for helpful discussions. This work was funded by the National Institute of General Medical Sciences R01 GM126112, and Silicon Valley Community Foundation, Chan Zuckerberg Initiative Grant DAF2021-238797, both to S.A.M. S.A.M. is supported by an Allen Distinguished Investigator Award (through the Paul G. Allen Frontiers Group), a Vallee Scholar Award, a Sloan Research Fellowship, and a New York Stem Cell Foundation Robertson Investigator Award; K.K. is supported by a Japan Society for the Promotion of Science Postdoctoral Fellowship; N.Y. is supported by McDonnell International Scholars Academy fellowship.

## Author Contributions

Conceptualization, Methodology, K.J., S.A.M.; Software, K.J.; Formal Analysis, K.J., N.Y.; Experimentation, K.J., M.T.A., H.W., X.Y., S.A.M., K.K., G.R.G.; Data Curation, K.J., M.T.A.; Writing – Original Draft, K.J., S.A.M.; Writing – Review & Editing, K.J., M.T.A., H.W., N.Y., K.K., G.R.G., S.A.M.; Visualization, K.J., S.A.M.; Funding Acquisition, Resources, Supervision, S.A.M.

## Methods

### Tissue culture

#### Isolation of mouse LSK cells

Lin^-^ Sca1^+^ c-Kit^+^ (LSK) cells were obtained using a previously described protocol^13^. Adult mice were euthanized, bone marrow was extracted from long bone, hips and spine and passed through a 70µm filter. Cells were centrifuged at 300g for 10mins at 4C and the pellet was resuspended in EasySep buffer (STEMCELL, Cat. 20144) at 100 million cells/ml. EasySep lineage depletion kit (STEMCELL, Cat. 19856) was used to remove differentiated cells. Finally, cells were stained for Sca1 (Sca1-AF488; BioLegend clone D7) and cKit (CD117-PE; BioLegend clone 2B8) and sorted using the MoFlo Cell Sorter (Beckman Coulter) with a 130µm nozzle. Isolated LSK cells were counted and used directly for lineage tracing experiments.

#### Mice and derivation of mouse embryonic fibroblasts

MEFs were derived from embryonic day (E)13.5 C57BL/6J embryos. (The Jackson laboratory: 000664). Heads and visceral organs were removed and the remaining tissue was minced with a razor blade and then dissociated in a mixture of 0.05% trypsin and 0.25% collagenase IV (Life Technologies) at 37 °C for 15 min. After passing the cell slurry through a 70-μM filter to remove debris, cells were washed and then plated on 0.1% gelatin-coated plates, in DMEM supplemented with 10% FBS (Gibco), 2mM l-glutamine and 50mM β-mercaptoethanol (Life Technologies). All animal procedures were based on animal care guidelines approved by the Institutional Animal Care and Use Committee.

### General Experimental methods

#### Lenti- and retro-virus production

Lentiviral particles were produced by transfecting 293T-17 cells (ATCC: CRL-11268) with the pSMAL-CellTag construct (see below), along with packaging constructs pCMV-dR8.2 dvpr (Addgene plasmid 8455), and pCMV-VSVG (Addgene plasmid 8454). Constructs were titered by serial dilution on 293T cells. Hnf4α -T2A-Foxa1 was cloned into the pGCDN-Sam retroviral construct and packaged with pCL-Eco (Novus Biologicals, NBP2-29540), titered on fibroblasts. We opted to generate a bicistronic *Hnf4α-Foxa1* construct, based on the T2A sequence to increase the consistency of reprogramming via maintenance of exogenous transcription factor stoichiometry. Virus was collected 48 h and 72 h after transfection and applied to cells immediately following filtering through a low-protein binding 0.45-μm filter. Wherever applicable, the virus was concentrated using high-speed centrifugation. 20ml of filtered viral supernatant was centrifuged at 50,000g for 2.5 hours at 4°C, supernatant was removed and the virus was resuspended in 100ul of DMEM. The concentrated virus was stored at -80C.

#### scRNA-seq library preparation

3’ single-cell RNA library preparation was performed using the Chromium Single Cell Gene Expression Kit from 10x Genomics. Cells were obtained as single-cell suspensions and processed according to the manufacturer’s instructions (CG000315).

#### CellTag amplification for scRNA-seq (CellTag-RNA PCR)

An additional PCR step was used to amplify CellTag barcodes from the single-cell cDNA library, obtained after step 2.4 of the 10x Genomics Single Cell Gene Expression Kit user guide (CG000315). 5ul (or at least 60ng) of cDNA was mixed with 2x Q5 HF PCR Master Mix (New England Biolabs) and 500nM of *P5/R1-par* and *P7/SI-R2* primers in a 50ul reaction volume and subjected to the following PCR program: 98 C for 30 seconds; N cycles (98°C for 10 seconds; 54°C for 30 seconds; 72°C for 30 seconds); 72°C for 2 minutes. The number of PCR cycles (N) was kept the same as the number of cycles used during sample index PCR of the main scRNA-seq library. CellTag amplicon library was purified using double-sided bead purification (0.4x-0.64x) and quantified on an Agilent TapeStation using the D1000-HS tape. Libraries were either sequenced by themselves (with a 50% Phi-X spike-in) or along with scRNA-seq libraries (preferred). CellTag amplicon libraries were sequenced on an Illumina NextSeq-500 to avoid index hopping-related artifacts. Primer sequences are available in Supplementary Table 1.

#### scATAC-seq library preparation

Standard scATAC-seq library preparation was performed using the Chromium Single Cell ATAC Kit from 10x Genomics. Cells were obtained as single-cell suspensions, nuclei were isolated using 10x Genomics nuclei isolation protocol (CG000169), and libraries were prepared according to the manufacturer’s instructions (CG000209).

#### scATAC-seq library preparation with modifications for CellTag capture

To prepare single-cell ATAC libraries with CellTag capture, nuclei were isolated using manufacturer’s instructions (CG000169), centrifuged to remove supernatant, and lightly fixed in 100ul 0.1% formaldehyde solution for 5 minutes. The reaction was stopped for 5 minutes by adding 30ul of stop buffer (0.625M Glycine, 0.5% BSA, 0.25M ph8 Tris-Cl in PBS). The nuclei suspension was diluted using 100ul diluted nuclei buffer (10x Genomics; CG000169) and pelleted using centrifugation. The pellet was subjected to tagmentation for 60 minutes after re-suspension in a 15ul tagmentation reaction (for up to 15k nuclei) according to the manufacturer’s instructions (CG000209). After tagmentation, the reaction mixture was diluted with 100ul dilute nuclei buffer, nuclei were pelleted using centrifugation and subjected to targeted in situ reverse transcription in a 100ul reaction volume (20ul of 5x SuperScript IV reaction buffer, 5ul each of dNTPs, DTT, RnaseOUT RNase inhibitor, SuperScript IV Reverse Transcriptase, 1uM of primer *ctac2-rt1*) using the following temperature program: 4°C for 2 minutes; 10°C for 2 minutes; 20°C for 2 minutes; 30°C for 2 minutes; 40°C for 2 minutes; 45°C for 10 minutes. After isRT, the reaction mixture was diluted with 100ul dilute nuclei buffer and pelleted using centrifugation. 15ul GEM-nuclei mix was prepared to load nuclei on 10x Genomics Chip E/H by mixing up to 15k nuclei with 6ul of ATAC buffer (from the 10x Genomics scATAC-seq kit) and 3ul of 4uM primer *ctac2-rt1*. Any remaining volume was made up with dilute nuclei buffer. GEM-nuclei mix was loaded onto Chip E/H along with ATAC GEM beads and barcoding enzyme mix, the remaining steps of the scATAC-seq library preparation protocol were performed according to the manufacturer’s instructions. Primer sequences are available in Supplementary Table 1. All centrifugation steps were performed at 500g for 10 minutes at 4°C unless stated otherwise.

#### CellTag amplification for scATAC-seq (CellTag-ATAC PCR)

While CellTags can be recovered directly from the sequenced scATAC-seq library with our library preparation, a higher yield can be obtained using an additional targeted PCR step, similar to the scRNA-seq version. For this, 5ul of the library is collected after step 3.2 of the user guide (CG000209) and mixed with 2x Q5 HF master mix, 500nM of primer *biot-atac2_lin* and water in a 50ul reaction volume, and CellTag containing fragments are linearly amplified using the following PCR program: 98°C for 30 seconds; 20 cycles (98°C for 10 seconds; 67°C for 30 seconds; 72°C for 30 seconds); 72°C for 2 minutes. The CellTag amplicons are purified using streptavidin-coated magnetic bead pulldown (ThermoFisher Scientific; Dynabeads™ MyOne™ Streptavidin C1) and purified fragments are resuspended in 20ul of water. A final sample index PCR is performed to create a sequencible library in presence of 2x Q5 master mix, 500nM each of *partial_p5* and *biot-atac2_e-rev* primers in a 100ul reaction volume using the following PCR program: 98°C for 30 seconds; 13 cycles (98°C for 10 seconds; 67°C for 30 seconds; 72°C for 30 seconds); 72°C for 2 minutes and libraries are purified using a double-sided bead cleanup, as described in Step 4.2 of 10x Genomic scATAC-seq user guide (CG000209). Primer sequences are available in Supplementary Table 1.

### General Computational methods

#### Identifying clones

Clone identification was performed based on our previously described method^7, 17^. Reads matching the CellTag-multi barcode sequence pattern (N)_3_GT(N)_3_CT(N)_3_AG(N)_3_TG(N)_3_CA(N)_3_ were extracted from single-cell bam files as obtained from CellRanger, filtered to remove false positive transcriptomic/genomic reads and reads originating from non-cell droplets. For scRNA-seq, cell barcode-CellTag-UMI triplets represented by only a single read were discarded. We also provide an estimate of CellTag sequencing saturation to guide users if they require deeper sequencing of their CellTag libraries. CellTags were error-corrected using Starcode^75^ to mitigate PCR/sequencing errors and filtered to remove sequences outside of the allowlist. Cell x CellTag read count (ATAC)/ UMI count (RNA) matrices were obtained, binarized and cells with too few or too many tags were removed to obtain the final Cell x CellTag matrices for scRNA-seq and scATAC-seq assays. Cell-cell similarity was computed using the Jaccard similarity metric and clones were identified using graph clustering. Whenever applicable, scRNA-seq and scATAC-seq CellTag matrices were merged before the Jaccard similarity calculation step, to identify clones across single-cell modalities. A detailed pipeline for clone calling can be found at: https://github.com/morris-lab/newCloneCalling

#### Clone cell embedding

To jointly visualize cells and clones on a single embedding, we developed a unique clone-cell graph embedding approach wherein we impute a cell-cell similarity graph with abstract clone nodes and use it as an input for graph embedding algorithms such as UMAP. For clone-cell embedding, we first obtained our single-cell data as an AnnData object and computed a cell-cell connectivity matrix based on PCA (in case of scRNA-seq) or CCA (in case of joint scRNA-seq scATAC-seq embedding). Next, we created a new AnnData object containing both cells and clones as observations. The connectivity matrix in the .obsm[‘connectivities’] slot was expanded to introduce clones. Then, clones were connected to their constituent cells by setting the respective entries in the expanded ‘connectivities’ matrix to 1. Finally, we used this clone-cell AnnData object with the expanded connectivity matrix as an input to graph embedding algorithms such as UMAP or Force Atlas.

### Section 1

#### CellTag-multi library synthesis

CellTag-multi library was synthesized using Restriction Free (RF) cloning^76^. CellTag-multi barcodes were obtained as a gBlock from IDT DNA (see Supplementary Table 1 for sequence) and cloned into the pSMAL-ctac2 vector. 20ng of the CellTag-multi-v1 gBlock and 100ng of pSMAL-ctac2 vector were mixed with 2x Phusion PCR master mix in a 20ul reaction volume. The reaction mixture was subjected to the following thermal cycling program: 98°C for 30 seconds; 15 cycles (98°C for 8 seconds, 60°C for 20 seconds, and 72°C for 4.5 minutes); 72°C for 5 minutes. The parental plasmid was digested by adding 2ul of methylation-sensitive restriction enzyme, *DpnI* (New England Biolabs), and incubating the reaction at 37°C for 2 hours followed by inactivation at 80°C for 20 minutes. 10ul of the reaction mix was transformed directly into 100ul of Stellar chemically competent cells (Takara Bio), cells were allowed to recover at 37°C, 250rpm in 1ml of SOC media and plated on a Nunc Square BioAssay plate (Cat. 166508). Plates were incubated overnight at 37°C. Bacterial colonies were collected using a scraper and allowed to recover in 150ml of LB media supplemented with 100ug/ml Ampicillin. CellTag-multi libraries were purified using a Qiagen High speed maxi prep kit (Cat. 12662) and library complexity was assessed as described below. This cloning was performed four times and libraries from each round were pooled to obtain the final high complexity library.

#### Assessing the complexity of CellTag-multi libraries and allowlisting

A list of allowed CellTag sequences for each CellTag library was created using amplicon sequencing. 50ng of CellTag plasmid library was mixed with 2x Q5 HF Master Mix, 2.5ul each of 0.5uM primers *bATAC_fwd* and *bATAC_rev* in a 25ul reaction volume and subjected to the following PCR program: 98°C for 30 seconds; 10 cycles (98°C for 10 seconds; 63°C for 30 seconds; 72°C for 1 minute). Two amplicon libraries were generated from each CellTag library plasmid preparation in parallel and sequenced on an Illumina Miseq. For each replicate, reads matching the CellTag sequence pattern (N)_3_GT(N)_3_CT(N)_3_AG(N)_3_TG(N)_3_CA(N)_3_ were extracted, sequencing/PCR errors were corrected by collapsing tags within 4 edits of each other using starcode^75^ and thresholded to retain CellTags containing at least N reads where N = max(10, 90^th^ percentile/10). An allowlist was created by collecting all CellTag sequences retained in thresholded lists from both replicates. Allowlists from the four CellTag libraries were combined to create the master allowlist for the CellTag-multi library (Supplementary Table 7). The detailed analysis code can be found at: https://github.com/morris-lab/newCloneCalling

#### Species mixing experiment

For the species mixing experiment, mouse iEP-LT cells were tagged with CellTag-multi-v1 library, containing the barcode pattern (N)_3_GT(N)_3_CT(N)_3_AG(N)_3_TG(N)_3_CA(N)_3_ and human HEK 293T cells with CellTag-multi-v0 library, containing the barcode pattern (N)_5_GTA(N)_5_CCT(N)_5_ATC(N)_5_GAT(N)_5_. Nuclei were isolated from both species using the Nuclei Isolation for scATAC-seq protocol from 10x Genomics (CG000169) and mixed in a 1:1 ratio. The mixed nuclei sample was processed using the standard scATAC-seq library preparation protocol (v1 kit) from 10x Genomics with modifications to capture CellTags. Single-cell libraries were sequenced on an Illumina Nextseq-500. The resulting sequencing data was aligned to a mixed species reference using CellTag-ATAC v1. The aligned bam file was used for downstream analysis.

Reads matching v0 or v1 CellTags were parsed from the mixed species single-cell aligned bam file. Each cell barcode was assigned to one of four categories, based on CellRanger-ATAC species assignments - human, mouse, doublet, non-cell; the distribution of v0 and v1 reads was assessed across the four categories. Cells with fewer than two CellTag reads across both libraries were discarded and the remaining cells were plotted on a barnyard plot. We quantified inter-species cross-talk of CellTags, by calculating the percent of cells, with at least 2 CellTag reads/cell, having less than 95% of CellTag reads originating from the correct, species-specific CellTag library.

#### Assessing the effect of isRT on chromatin accessibility signal

We compared the effect of introducing an isRT step on scATAC-seq data quality. For this, two single-cell ATAC libraries were prepared with CellTagged HEK 293T cells using either the original 10x Genomics scATAC library preparation protocol (Original) or our modified method (Modified). Sequencing data from both was processed with ArchR^77^, dimensionally reduced using LSI, clustered using Louvain clustering, and peaks were identified across samples. Both datasets were compared across several standard scATAC-seq data quality metrics such as fragment size distribution, TSS scores, the number of unique fragments per cell and Fraction reads in Peaks (FRiP) per cell. To compare genome-wide accessibility data across samples, normalized peak counts (Counts Per Million; CPM) were calculated for each sample and plotted on a scatter plot and the Pearson Correlation coefficient was calculated to quantify the similarity between the accessibility signal of the two samples.

#### Analysis of clones in expanded reprogramming fibroblasts

A subset of the data obtained from our reprogramming dataset (described in section 3) from Days 12 and 21 was used for this analysis. Clones were identified following the standard computational workflow as described above. CellTag abundance was calculated for each CellTag as the percent of metric filtered cells containing that CellTag. Browser tracks depicting single-cell accessibility fragments were plotted using ArchR. Gene expression and gene scores values were averaged on a clonal level. Spearman correlation coefficients were calculated between clonal gene expression and gene score both within (Intra clonal) and across clones (Inter clonal).

### Section 2

#### Lineage tracing during in vitro mouse hematopoiesis

LSK cells were purified as described above, counted and 5,500 cells were added to a 96-well U-bottom suspension culture plate (GenClone Cat. 25-224) and allowed to recover in broad myeloid differentiation media^13^ consisting of SFEM media (STEMCELL), Pen/Strep, IL-3 (20ng/mL; PeproTech Cat. 213-13), FLT3-L (50ng/mL; PeproTech Cat. 250-31L), IL-11 (50ng/mL; PeproTech Cat. 220-11), IL-5 (10ng/mL; PeproTech Cat. 215-15), EPO (3U/mL; PeproTech Cat. 100-64), TPO (50ng/mL; PeproTech Cat. 315-14), and mSCF (50ng/mL; R&D Systems Cat. Q78ED8) and IL-6 (10ng/mL; R&D Systems Cat. 406-ML-005) at 37°C for 2 hours.

To allow clone tracking, cells were transduced for 2 days with 10ul of concentrated CellTag-multi virus (∼25k unique CellTag sequences) in 100ul differentiation media, in the presence of 6ug/ml DEAE-Dextran after spin-fection at 800g for 90 minutes at 37°C. 60 hours (2.5 days) after the start of the experiment, 50% of the cells were collected for single-cell profiling and split equally between scRNA-seq and scATAC-seq assays. The remaining cells were split into 2 technical replicates and re-plated in fresh differentiation media. Finally, all the cells were collected on Day 5 and split between scRNA-seq and scATAC-seq profiling.

#### Single-cell library preparation and sequencing

The v3 single index Gene Expression kit and the v1 scATAC kit from 10x Genomics were used for single-cell library preparation. CellTag-RNA PCR was used to obtain CellTag amplicon libraries as described above. scRNA-seq libraries were sequenced on an Illumina NovaSeq-6000 and the resulting data was computationally dehopped. CellTag amplicon libraries obtained from scRNA-seq libraries were sequenced on an Illumina NextSeq-500. For read alignment, CellTag and transcriptome read files for each sample were processed together using CellRanger, using a custom mm10 reference containing the GFP CDS and UTR, to produce one aligned bam file per sample. scATAC-seq libraries containing both accessible chromatin and CellTag fragments were sequenced on an Illumina NextSeq-500 and processed using CellRanger-ATAC, using the default mm10 reference genome. Aligned bam files from both modalities were used for CellTag processing, other CellRanger and CellRanger-ATAC outputs were used for downstream single-cell analyses.

#### Basic single-cell and clonal analysis of the Hematopoiesis dataset

CellRanger generated scRNA-seq count matrices were processed using Seurat. Low-quality cells with high mitochondrial reads, low UMIs, and features per cell were removed. Day 2.5 and Day 5 samples were integrated using SCTransform, dimensionally reduced using PCA, and clustered using Louvain clustering. scRNA-seq clusters were annotated using marker gene expression. Fragments files from scATAC-seq samples were processed using ArchR v1.0.1. Valid cell barcodes, as identified by CellRanger-ATAC and passing default ArchR quality filters were retained. Cells were dimensionally reduced using iterative LSI and clustered using Louvain clustering. Cell type labels were transferred to scATAC-seq clusters using Seurat label transfer and annotations were manually inspected using marker gene scores. For RNA-ATAC co-embedding, scRNA-seq gene expression matrix and scATAC-seq MAGIC imputed^78^ Gene Score matrix, as obtained from ArchR, were used as input to the RunCCA function in Seurat. A union set of the top 5000 highly variable genes from each dataset were used for this co-embedding.

For clone calling, reads mapping to the CellTag barcode were extracted from single-cell aligned bam files as obtained from CellRanger and CellRanger-ATAC and cell x CellTag UMI matrices were obtained. CellTag data within each modality was merged, retaining sample-of-origin information in the cell barcode, and cell x CellTag UMI (for RNA) and read (for ATAC) count matrices were obtained for each modality. The RNA matrix was binarized at a threshold of more than one UMI count and cells with 2 to 25 CellTags were retained. The ATAC matrix was binarized at a threshold of more than one read count and cells with 1 to 25 CellTags were retained. The two filtered matrices were merged, cell-cell Jaccard similarity matrix was computed and thresholded at 0.6 (for cell pairs within the same modality) and 0.5 (for cell pairs across modalities). The final thresholded matrix was used to identify clones across the entire dataset. Clone-cell embedding was computed as described above, and ForceAtlas2 was used to jointly visualize clones and cells. This embedding was also generated separately for sub-clones where clones were split either by modality or by both, time point and modality. For single-modality clonal analysis, Cell x CellTag matrices for each modality were processed separately with the same thresholds as above. A Jaccard threshold of 0.5 was used for ATAC clone calling and 0.6 was used for RNA clone calling. Lineage hierarchies were obtained using clone coupling as previously described^13^

#### State-fate linkage in hematopoiesis

To link cell state with fate, we first obtained all clones spanning the two time points (state-fate clones). Each state-fate clone was assigned a fate label, which was the most common fate amongst its Day 5 sisters. Less common lineages were grouped based on similarity, e.g. Erythroid and Megakaryocytes (Ery/Meg); Eosinophils, Basophils, and Mast Cells (Baso/Eos/Mast). Ccr7 DCs and plastoid DCs (DCs). Clones annotated to transition/ progenitor fates were excluded from state-fate analysis unless otherwise specified. Fate bias scores were calculated as percent of Day 5 fate sisters belonging to the annotated fate label.

To map Day 2.5 (state) sub-clones on the clone-cell embedding, we split each clone into sub-clones based on the time point of collection and assay of each sister, to obtain up to four sub-clones RNA/ATAC – state/fate sub-clones. The clone-cell embedding was recomputed using these sub-clones. Overlap between RNA and ATAC sub-clones across the two single-cell modalities was calculated within each ‘fate potential’ group using the Wasserstein distance metric computed with a 30-dimensional embedding of the sub-clone nodes obtained using the UMAP algorithm. To quantify if state sub-clones closer to the periphery of a ’fate potential’ group were less fate biased, we devised a closeness metric, which is the minimum distance of a state sub-clone from the centroid of an alternative fate potential group. A higher closeness metric would mean that a state sub-clone is farther away from centroids of other fate potential groups. The relationship between the closeness metric and fate bias was plotted using a percentile plot, with percentile rank for the closeness metric on the X-axis and mean fate bias scores for state sub-clones passing the percentile rank on the Y-axis.

To characterize functional priming of cell state, Day 2.5 state sisters in each fate potential group were compared to the rest in gene expression and TF activity space. For scRNA-seq features, we used residuals obtained for the top 3000 highly variable genes after SCTransform normalization in Seurat. For scATAC-seq features, we used TF activity z-scores obtained from chromVAR using the default mouse motif set in ArchR (884 TF motifs). Correction for multiple hypothesis testing was performed using the Benjamini-Hochberg method, setting the FDR threshold for significance at 0.05, unless otherwise specified.

#### Fate prediction from cell state using machine learning

State-fate machine learning was performed to quantify the predictability of cell fate from early state. A machine learning classifier was tasked to predict the discrete clonal fate label Y as obtained above (possible fate labels: ‘progenitor’, ‘monocyte’, ‘neutrophil’, ‘Lym/pDC/Ccr7-DC’, ‘Ery/Meg’ or ‘Baso/Eos/Mast’), from an input vector of single-cell features X of Day 2.5 cells. For RNA only model, we used residuals of the top 3000 genes for input, for ATAC only model, we used TF activity z-scores (with k-nn imputation where k=20) as input and for the RNA+ATAC model, we randomly paired RNA and ATAC cells within the same sub-clone and concatenated their respective RNA and ATAC feature vectors and used those as input. For training, we used the Repeated Stratified k-fold cross-validation procedure setting both *n_splits* and *n_repeats* to 5. Model performance was evaluated using accuracy and Weighted F1 score.

For each machine learning task, we tested a panel of classifier architectures, logistic regression, LightGBM, and Random Forest. Each was trained and evaluated using the procedure described above. Hyperparameter tuning was performed for each and the following values were tested:

- Random Forest: n_estimators: [100, 300, 1000], max_depth: [10, 50, None], min_samples_leaf: [1, 2, 4], bootstrap: [True, False]
- LightGBM: num_leaves: [7,15,31,80], max_depth: [5,9,30], min_data_in_leaf: [20, 40, 80], bagging_fraction: [0.8,1], bagging_freq: [3], feature_fraction: [0.1, 0.9]
- Logistic Regression: penalty: [’l2’, ’none’], C: np.logspace(−4, 4, 20), solver: [’lbfgs’,’newton-cg’,’saga’], max_iter: [1000]

The python library ‘scikit-learn’ was used for all machine learning analysis.

#### Fate prediction using TF activities derived from distal, intronic, exonic, and promoter peak sets

ATAC peaks were categorized as intronic, exonic, promoter, or distal using default ArchR definitions. TF activity scores were calculated for each peak set independently and used for state-fate prediction analysis as described above. To test if variation in model performance was due to different numbers of peaks in each set, all peak sets were randomly sub-sampled to 8823 peaks (number of peaks in the exonic set), TF activity scores were calculated again and state-fate prediction was performed using these new scores.

#### SHapley Additive exPlanations (SHAP) analysis

The shap python package was used to perform SHAP analysis and interpret trained machine learning models. The TreeExplainer function from the ‘shap’ python package was used to calculate SHAP values for trained random forest models. For each input feature and fate label, SHAP values were calculated using each data point in the 25 test sets (*n_splits* x *n_repeats*), resulting in 5 SHAP values per data point per feature. This helped average out any rare outlier values generated due to a model training artifact. Feature importance scores were calculated for each input feature for the prediction of each fate label, by taking the mean of absolute SHAP values for each feature-fate combination. To identify features positively or negatively correlated with the prediction of a fate label, SHAP correlation was performed. For each input feature, the Pearson correlation coefficient between its values (expression/TF activity) and its SHAP values for a given fate was calculated, resulting in one correlation value per feature per fate.

### Section 3

#### Lineage tracing during iEP reprogramming

Cryo-preserved P0 MEFs were thawed and seeded in 0.1% gelatin-coated six-well plates, in DMEM supplemented with 10% FBS, 2 mM l-glutamine, and 50 mM β-mercaptoethanol (Life Technologies) and penicillin-streptomycin at a density of 30,000 cells/well. After overnight recovery at 37°C, cells were transduced every 12 hours for 2 days, with fresh Hnf4α-T2A-Foxa1 retrovirus in the presence of 4 μg/ml protamine sulfate (Sigma-Aldrich). During the last round of transduction, the retroviral mix was supplemented with CellTag-multi lentiviral library to initiate clone tracking. On Day 0 of reprogramming, cell culture media was changed to hepato-medium (DMEM:F-12, supplemented with 10% FBS, 1 μg/ml insulin (Sigma-Aldrich), 100 nM dexamethasone (Sigma-Aldrich), 10 mM nicotinamide (Sigma-Aldrich), 2 mM l-glutamine, 50 mM β-mercaptoethanol (Life Technologies), and penicillin-streptomycin, containing 20 ng/ml epidermal growth factor (Sigma-Aldrich)). After 72 hours (Day 3 of reprogramming), cells were dissociated, two-thirds of the cells were collected for single-cell sequencing and the remaining cells were re-plated on 6-well plates coated with 5 μg/cm^2^ Type I rat collagen (Gibco, A1048301). Two additional samples were collected on Days 11 and 21 for single-cell sequencing. We used the 10x Genomics v3.1 dual index Gene Expression kit (PN-1000268) and the v1.1 ATAC kit (PN-1000175) for single-cell profiling. This experiment was performed in two biological replicates.

CellTag PCR was performed for all scRNA-seq and scATAC-seq libraries, as described above. scRNA-seq and scATAC-seq libraries were sequenced on an Illumina NovaSeq-6000. CellTag amplicon libraries were sequenced on an Illumina NextSeq-500 to avoid any index hopping related artifacts. For read alignment, CellTag and transcriptome/chromatin read files for each sample were processed together using CellRanger/CellRanger-ATAC, to produce one aligned bam file per sample. Aligned bam files from both modalities were used for CellTag processing, other CellRanger and CellRanger-ATAC outputs were used for downstream single-cell analyses.

#### Basic single-cell and clonal analysis of the direct reprogramming dataset

scRNA-seq count matrices were processed using Seurat. Quality filtering was performed to remove cells with high mitochondrial reads and low UMIs and genes per cell. scRNA-seq samples across all time points and biological replicates were integrated, dimensionally reduced using PCA, and clustered using Louvain clustering. Cells from Days 12 and 21 were subsetted and re-clustered. Single-cell identity scores were obtained using Capybara, using Fibroblasts (MEFs), and reprogrammed, and dead-end trajectories from a previous dataset^7^ as references. Cell clusters were annotated as ‘reprogrammed’, ‘dead-end’, or ‘transition’ based on these cell identity scores and marker gene expression. Fragments files from scATAC-seq samples were processed using ArchR. Valid cell barcodes, as identified by CellRanger-ATAC and passing default ArchR quality filters were retained. Cells were dimensionally reduced using iterative LSI and clustered using Louvain clustering. Cells were annotated as ‘reprogrammed’, ‘dead-end’, or ‘transition’ based on marker gene accessibility. For RNA-ATAC co-embedding, scRNA-seq gene expression matrix and scATAC-seq MAGIC imputed^78^ Gene Score matrix, as obtained from ArchR, were used as input to the RunCCA function in Seurat. A union set of the top 2000 highly variable genes from each dataset were used for this co-embedding.

For clone calling, reads mapping to the CellTag barcode were extracted from single-cell aligned bam files as obtained from CellRanger and CellRanger-ATAC. CellTag data within each modality was merged, retaining sample-of-origin information in the cell barcode, and cell x CellTag UMI (for RNA) and read (for ATAC) count matrices were obtained for each modality. The RNA matrix was binarized at a threshold of more than one UMI count and cells with 2 to 25 CellTags were retained. The ATAC matrix was binarized at a threshold of more than one read count and cells with 2 to 25 CellTags were retained. The two filtered matrices were merged, cell-cell Jaccard similarity matrix was computed and thresholded at 0.6. The final thresholded matrix was used to identify clones across the entire dataset. Clone-cell embedding was computed as described above and the UMAP algorithm was used to jointly visualize clones and cells.

#### State-fate analysis for the direct reprogramming dataset

Clones were annotated with one of three fates – ‘reprogrammed’, ‘transition’, or ‘dead-end’, based on the most abundant cell type amongst fate sisters. Clonal fate bias scores were calculated as percent of fate sisters (Days 12 and 21) belonging to the annotated fate label. Alluvial plots were constructed using the ggAlluvial R package. State-fate machine learning analysis was performed exactly as described in the hematopoiesis section. Classification models were trained to predict either ‘reprogrammed’ or ‘dead-end’ fates. Since the frequency distribution of fate labels was less skewed for the reprogramming dataset, only prediction accuracy scores were used as performance metrics. CellRank analysis was performed for a 40,000-cell subset of the scRNA-seq dataset, due to scalability limitations. For feature enrichment analysis, Day 3 sisters in state-fate clones were grouped by fate. Seurat FindMarkers function was used to identify gene expression markers and ArchR getMarkerFeatures function was used to identify peak and TF activity markers for each of the following cell groups - uninduced MEFs, on-target destined cells and, off-target destined cells, in a series of one versus all comparisons. For peak and TF activity comparisons, both on-target and off-target cell groups were expanded using k-nearest neighbors (k=5). Uniquely enriched features (genes/peaks/TFs) were obtained by removing features that were identified as markers of more than one cell group. TF activity results were further refined by discarding TFs with low gene score-TF activity correlation (< 0.3). Motif enrichment analysis was performed using the HOMER package^79^ for both on-target and off-target DERs using MEF DERs as background, to better resolve fate-specific motif enrichment. Mouse ENCODE ELS elements were obtained from the ENCODE SCREEN database^43^. Only genomic regions annotated as dELS, pELS, dELS, CTCF-bound, or pELS, CTCF-bound in the SCREEN database were used for enrichment analysis. The FigR^42^ package was used for peak-to-gene linkage analysis. Optimal matching was used to pair RNA and ATAC cells from the same time points followed by the runGenePeakcorr function to identify peak-gene pairs. Peak-gene pairs with an adjusted p-value greater than or equal to 0.05 were discarded. *Foxa1* and *Hnf4α* ChIP-seq peaks from Day 2 of reprogramming were obtained^55^. These peak sets were added as custom annotations in ArchR and single-cell accessibility z-scores for each peak set were computed using the addDeviationsMatrix function in ArchR.

#### Computational analysis related to ZFP281 motifs

Tomtom analysis^58^ from the MEME-ChIP package was used to find highly similar motifs to *Zfp281*. The *Zfp281* position frequency matrix (PFM) was obtained from ArchR and used as input to the Tomtom web interface. Highly correlated TF motifs with q-value less than 0.05 were obtained, these were further subsetted for TF activities enriched in off-target destined cells resulting in a total of four TF motifs for comparison with *Zfp281*. *Zfp281* ChIP-seq peaks were obtained^56^ and single-cell accessibility z-scores were computed using the addDeviationsMatrix function in ArchR. *Zfp281* gene targets^59^ were used as inputs for a state-fate prediction model, which was trained and evaluated as described above and compared to a sized-matched set of random genes.

#### Plasmid cloning related to Foxd2 and Zfp281 experiments

Non-targeting shRNA construct was obtained from Sigma (SHC202; pLKO.5-puro Control Plasmid). Zfp281 targeting shRNA gene was obtained from Sigma (Clone ID: TRCN0000255746) and cloned into the pLKO.5-puro lentiviral construct (Sigma SHC201). For over-expression, cDNA fragments were cloned in the pGCDNsam retroviral construct. *Zfp281* cDNA was obtained from OriGene (Cat. MC205914) and *Foxd2* cDNA was reverse transcribed from RNA obtained from long-term iEP cells.

#### Reprogramming with Foxd2 and Zfp281 perturbations

Reprogramming was performed as described above, with the following modifications. For over-expression, cells were transduced with a 1:1 mixture of *Foxd2/Zfp281* retrovirus and *Hnf4α-Foxa1* reprogramming retrovirus every 12 hours for 2 days. Control cells were transduced with a 1:1 mixture of a GFP control retrovirus and *Hnf4α-Foxa1* reprogramming retrovirus for the same amount of time. For knockdown, cells were transduced with the non-targeting control/*Zfp281*-shRNA lentivirus every 12 hours for 1 day after the 2-day *Hnf4α-Foxa1* retroviral transduction was completed.

#### Single-cell analysis for Foxd2 and Zfp281 experiments

Dual indexed v3.1 scRNA-seq libraries were prepared for all four samples (*Zfp281* OE, OE Control, *Zfp281* KD, KD Control) according to the manufacturer’s instructions (CG000315) and sequenced on a Nextseq-500. Count matrices were generated and integrated using CellRanger count and aggr commands and processed using Seurat. Quality filtering was performed to remove cells with high mitochondrial reads and low UMIs and genes per cell. Cells were dimensionally reduced using PCA, cell cycle regressed, clustered using Louvain clustering, and visualized using UMAP. Capybara identity scores were calculated as described in the iEP lineage tracing section above. Markers for each lineage across time points and uninduced MEFs were obtained (log2 fold change > 0.7, adjusted p-value < 0.05) and used for gene module scoring for all four samples. Cell clusters displaying strong enrichment of on-target or off-target markers were annotated with the respective fates. pROGENY pathway analysis^62^ was used to calculate single-cell activity scores for the TGF-β signaling pathway.

#### Colony formation assays

Colony formation assays were performed as previously described^7^. Reprogramming cells were seeded at low plating density in collagen-coated 6-well plates within the first 4 days and allowed to form colonies over 2 weeks of reprogramming. Following this, cells were fixed using 4% paraformaldehyde, permeabilized using 0.1% Triton-X and processed for CDH1 (E-Cadherin) staining using the VIP peroxidase substrate kit (Vector labs SK4600) and anti-mouse E-Cadherin primary antibody (1:100, BD Biosciences). Stained colonies were imaged using a flatbed scanner and quantified using the following script: https://github.com/morris-lab/Colony-counter

#### Quantitative PCR and analysis

Cells were collected for RNA extraction (RNeasy kit; QIAgen) on Day 12 of reprogramming and reverse transcribed using the Maxima RT kit (ThermoFisher K1672). 20ng of reverse transcribed RNA was mixed with TaqMan™ Gene Expression Master Mix (ThermoFisher Scientific) and gene-specific TaqMan™ probes (Supplementary Table 8) in a 20ul reaction volume and processed according to manufacturer’s instructions (4371135) on the StepOne Plus qPCR system. Per gene fold change for Foxd2 overexpressing cells was calculated relative to control reprogramming cells (*Hnf4α*-*Foxa1* and GFP control overexpression) that were processed in parallel, after normalization to the housekeeping gene, *Actb*.

#### Reprogramming with TGF-β inhibition

Cells were reprogrammed as described above. Cells were cultured in hepatic media supplemented with 2.6µM of SB431542 (STEM CELL 72232), a small molecule inhibitor of TGF-b signaling starting on Day 0 of reprogramming. SB431542 containing media was changed every 2 days. Cells were collected for qPCR analysis on Day 5 of reprogramming and processed as described above.

