## Extended Figures for "Multiomic single-cell lineage tracing to dissect fate-specific gene regulatory programs"

#### Extended Figure 1

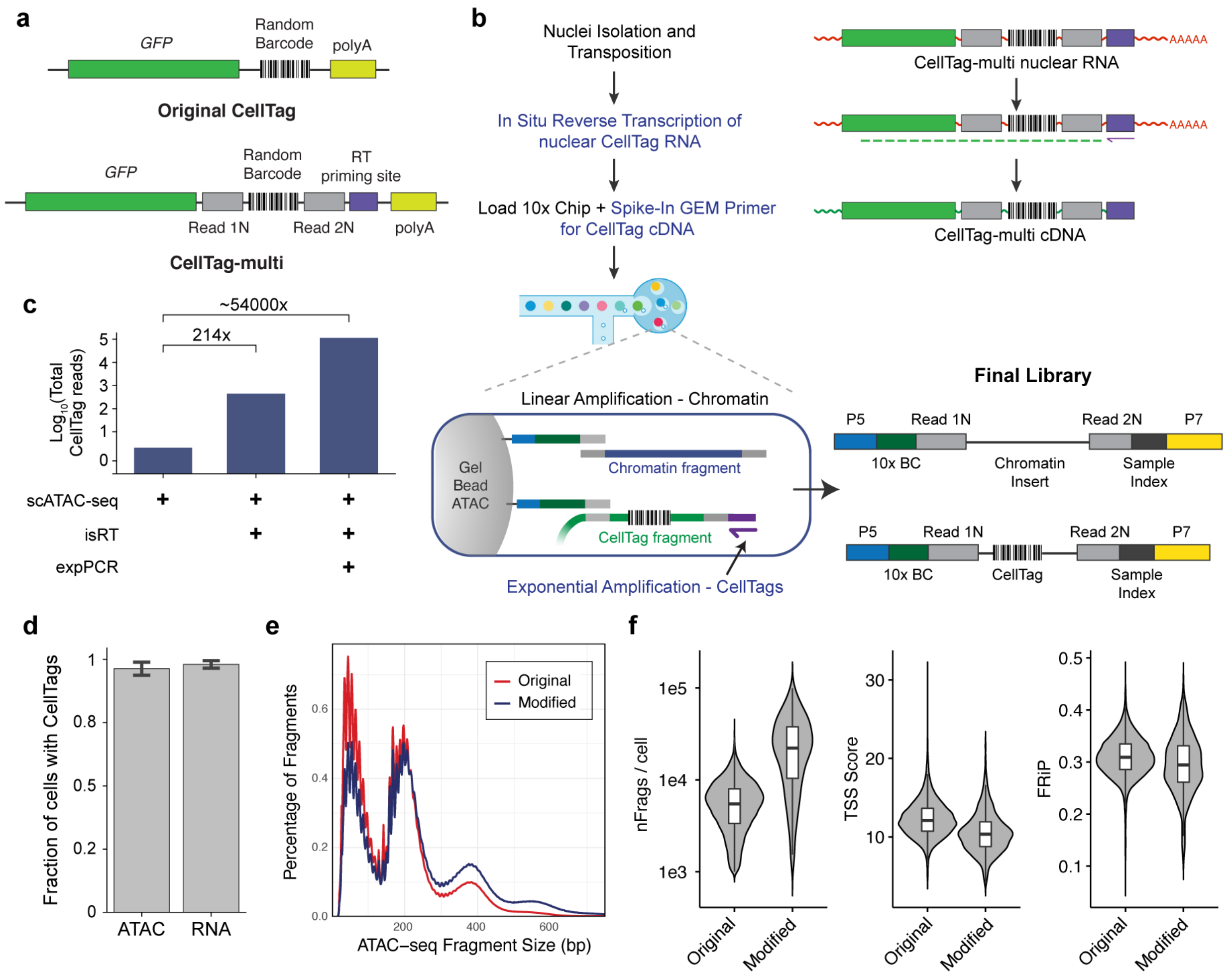


**Extended Figure 1. Development of CellTag-multi for parallel capture of lineage with scRNA-seq and scATAC-seq.** (**a**) Schematic comparing the original[7,17] CellTag lineage tracing construct to the new CellTag-multi construct. (**b**) Left Panel: Detailed flow chart and schematic of the modified scATAC-seq library preparation protocol. Right Panel: Major molecular steps of the protocol and the final library containing both CellTag and chromatin accessibility fragments. (**c**) Bar plot comparing total number of CellTag reads per library obtained across different scATAC-seq library preparation methods. Each library was sequenced to a similar sequencing depth. (**d**) Bar plot depicting fraction of cells with at least one CellTag detect across scRNA-seq and scATAC-seq samples. Percent cells with CellTags detected in scATAC-seq, relative to scRNA-seq. Comparison of (**e**) fragment size distribution and (**f**) various scATAC-seq quality metrics across two datasets generated using the original and modified scATAC-seq library preparation method (nFrags/cell: number of unique fragments per cell; FRiP: Fraction of reads in Peaks).

#### Extended Figure 2

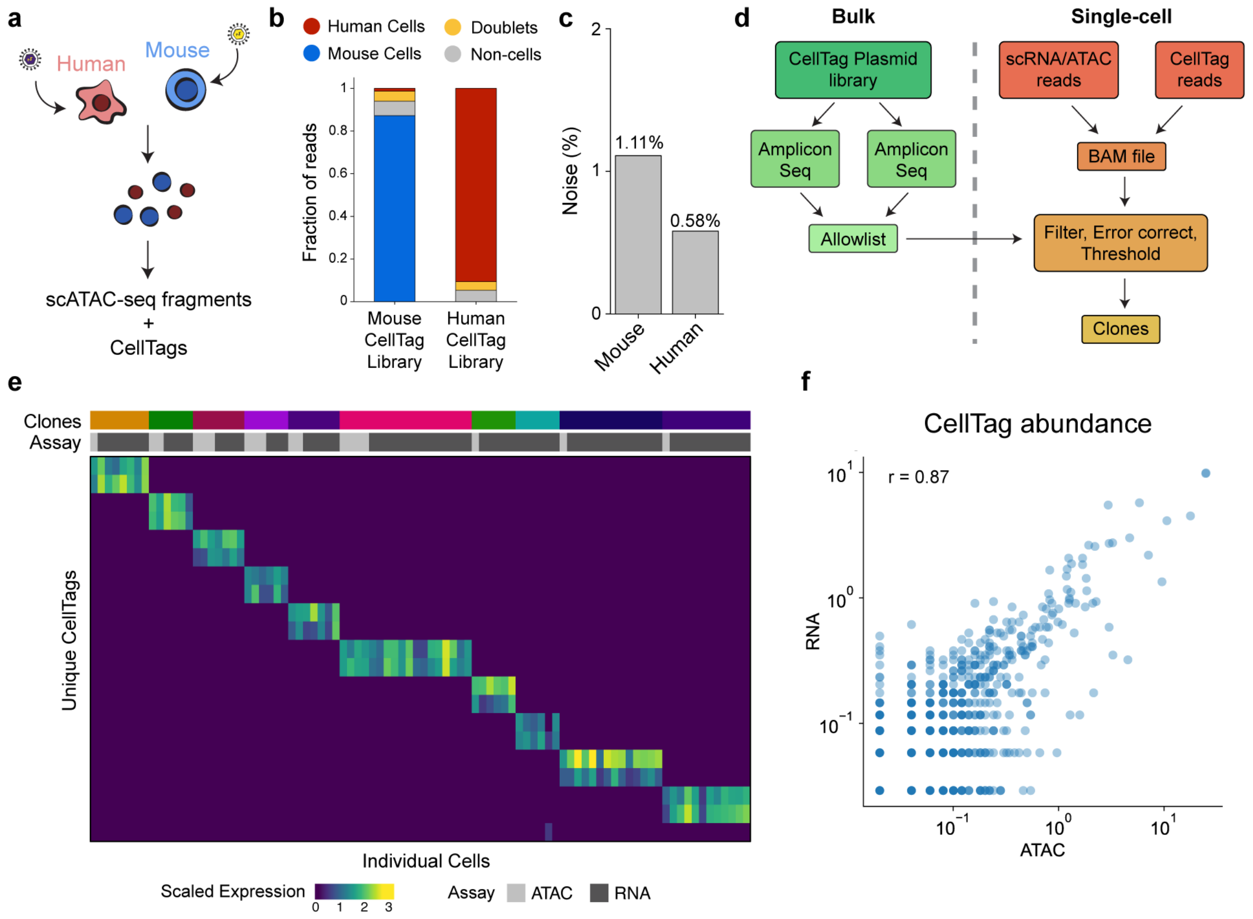


**Extended Figure 2. Testing CellTag-multi in cell lines and reprogramming fibroblasts.** (**a**) Schematic of the species mixing experiment to assess purity of CellTag signatures in scATAC-seq. (**b**) Bar plot depicting distribution of CellTag reads across, human, mouse, doublet and non-cell droplets for the two CellTag libraries. We observed that the majority of CellTag reads mapped to the expected species of origin, 87.2% for the mouse library and 91.4% for the human library. (**c**) To quantitively assess fidelity of CellTag signatures, we devised a measure for CellTag cross-talk rate (**Methods**). We find that for both human and mouse samples, the cross-talk levels were below 5%. (**d**) Schematic depicting the workflow for CellTag library allowlisting and clone identification from single-cell CellTag reads. (**e**) Heatmap depicting scaled CellTag expression across ten clones in a population of expanded reprogramming fibroblasts. (**f**) Correlation between CellTag abundance across scRNA-seq and scATAC-seq cells from the reprogramming dataset (Pearson's correlation coefficient = 0.87).

### Extended Figure 3

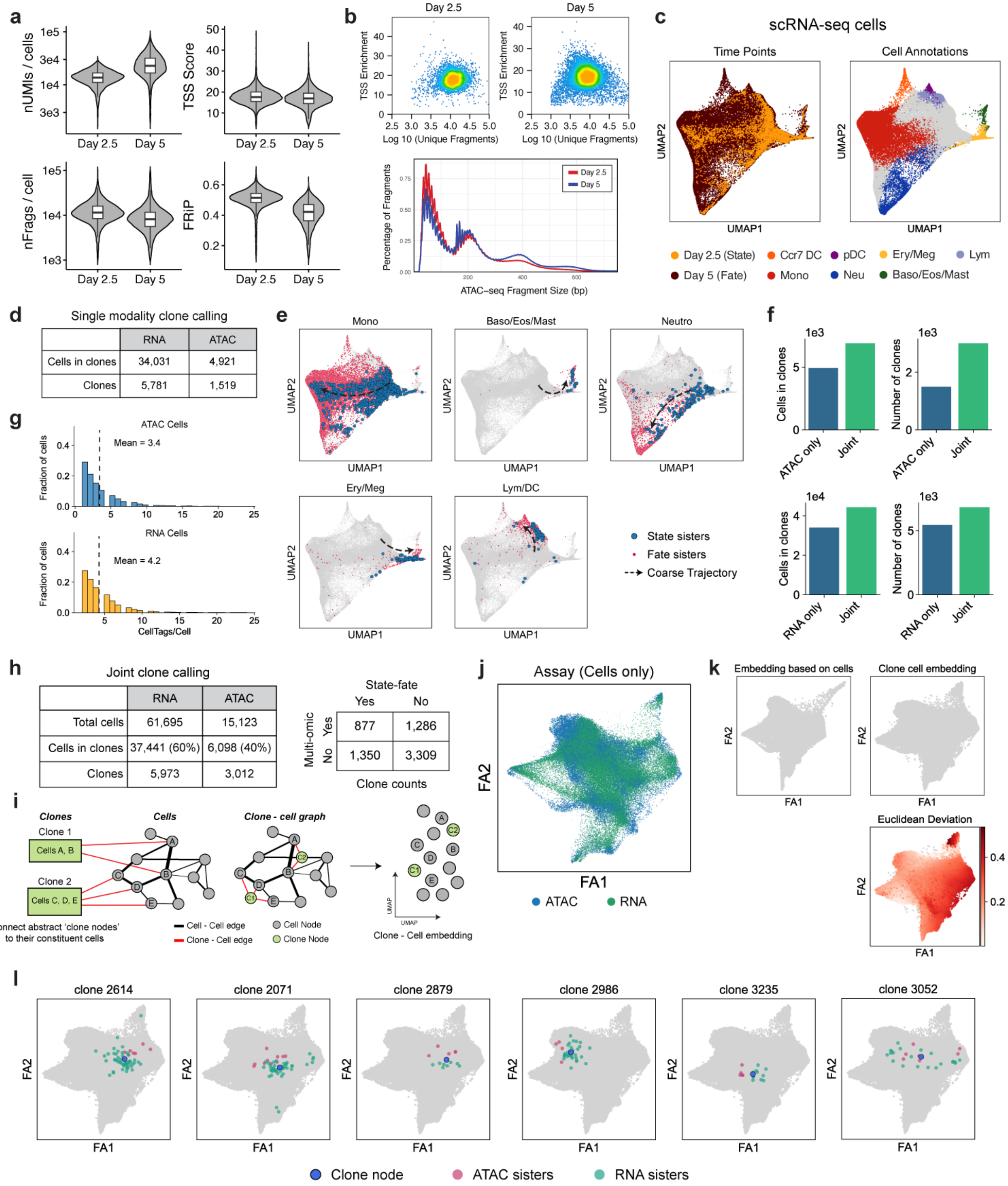

**Extended Figure 3. Single-cell metrics and cell annotation in hematopoiesis.** (**a**) Violin plots for single-cell quality metrics for the scRNA-seq and scATAC-seq datasets. (**b**) Unique fragments/cell vs single-cell TSS enrichment scatterplots and fragment size distribution plot for the two scATAC-seq time-points. (**c**) scRNA-seq UMAPs with time point (left panel) and cell fate information (right panel) projected. (**d**) Table summarizing clones identified in scRNA and scATAC datasets independently. (**e**) scRNA-seq UMAPs with state and fate sisters for major hematopoietic fates highlighted. (**f**) Bar plots comparing number of clones and cells in clones across single-modality and joint modality clone calling. (**g**) Histograms of CellTags detected per cell across scRNA-seq and scATAC-seq datasets after filtering and processing of CellTag reads. (**h**) Tables summarizing all clones identified in the dataset. (**i**) Workflow for joint embedding of cells and clone nodes. (**j**) Joint clone-cell graph based UMAP with assay information projected (only cells are shown). (**k**) Comparison of cell embeddings obtained using conventional UMAP vs a joint clone-cell graph based UMAP (only cell nodes shown, for direct comparison). Bottom Panel: Clone-cell graph UMAP with cells colored by deviation in their position on the UMAP between the two embeddings. (**l**) Visualization of clones along with their constituent cells confirms that clone nodes faithfully represent cells.

### Extended Figure 4

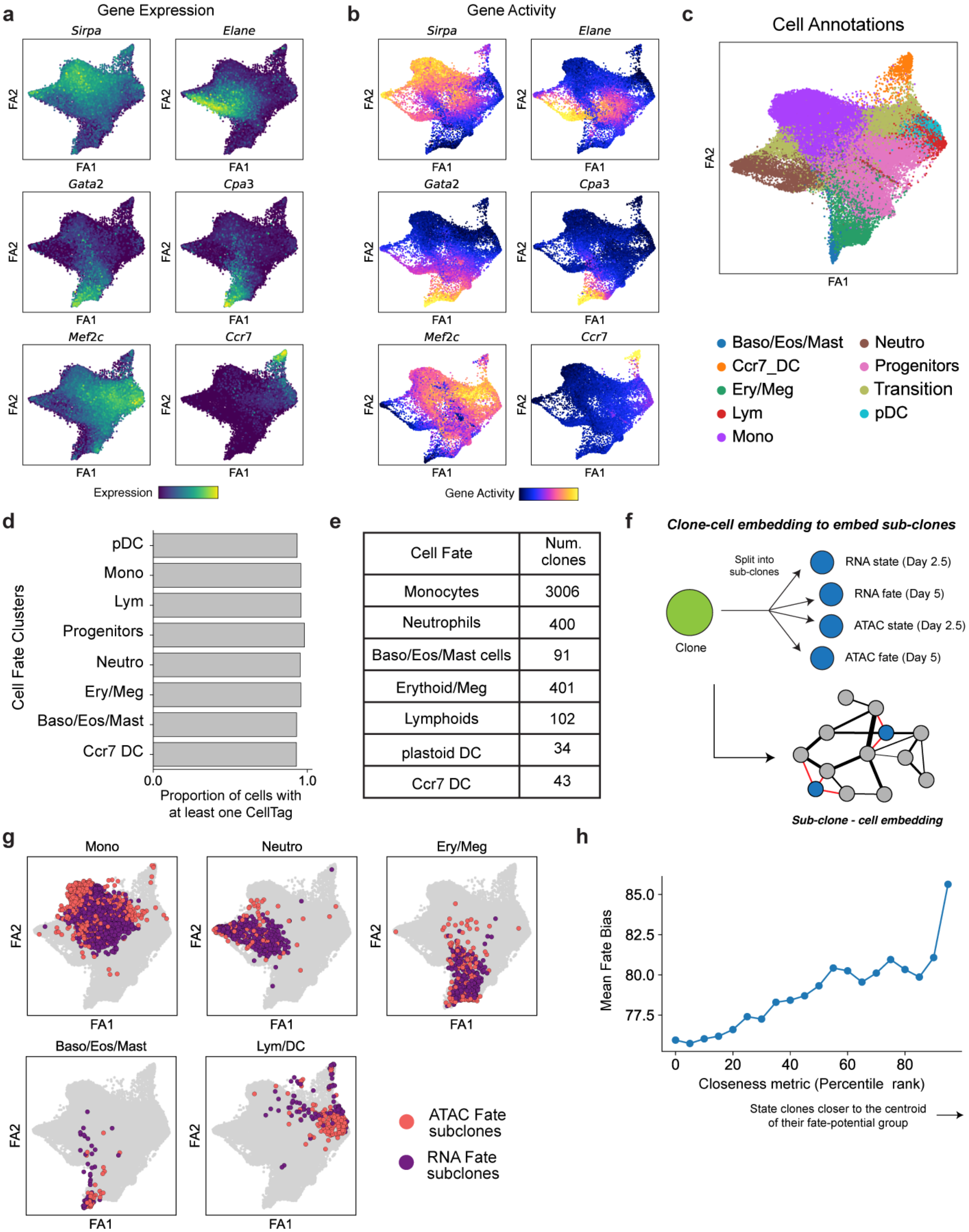


| Cell Fate | Num. clones |
|---|---|
| Monocytes | 3006 |
| Neutrophils | 400 |
| Baso/Eos/Mast cells | 91 |
| Erythoid/Meg | 401 |
| Lymphoids | 102 |
| plastoid DC | 34 |
| Ccr7 DC | 43 |

**Extended Figure 4. Fate annotation in hematopoiesis.** (**a**) Marker gene expression and (**b**) accessibility projected on the UMAP for various hematopoietic cell fates. (**c**) UMAP with the full set of cell annotations in the hematopoiesis dataset projected. (**d**) Bar plot summarizing proportion of cells with at least one detectable CellTag across major cell fate clusters. CellTags are profiled uniformly across all cell states. (**e**) Table summarizing number of clones identified in each fate. Clonal fate was annotated using the most dominant cell type amongst Day 5 fate sisters. (**f**) Schematic depicting joint embedding of sub-clones with cells using the clone-cell embedding method. (**g**) UMAP with fate sub-clone nodes for major lineages highlighted. (**h**) Plot showing that fate bias increases from the periphery of each state group towards the center. The closeness metric is directly proportional to the closeness of a state sub-clone node to the centroid of its state group in a 30-dimensional UMAP space (**Methods**).

#### Extended Figure 5

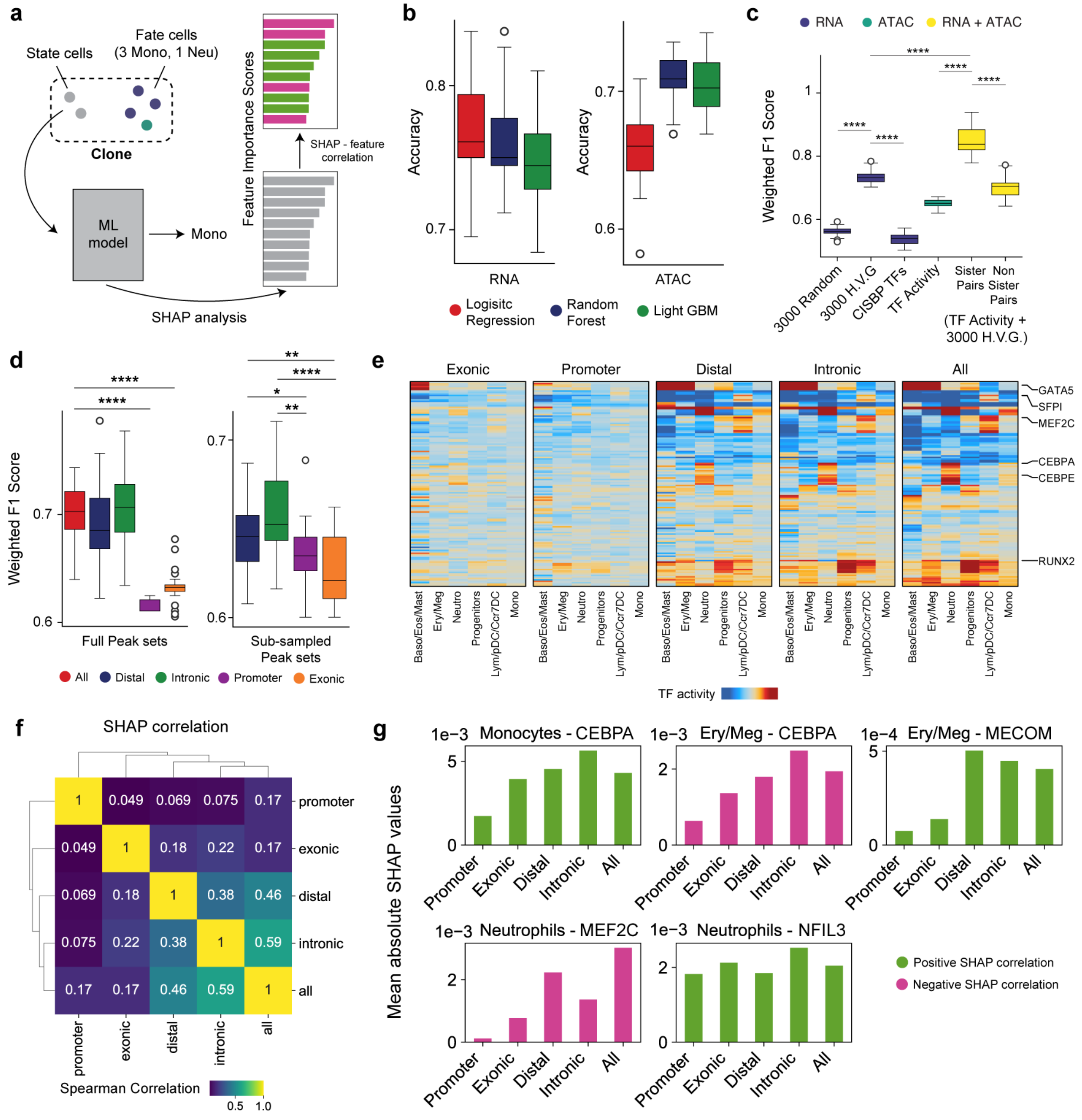


**Extended Figure 5. Machine learning analysis to predict cell fate from state.** (**a**) Schematic of state-fate prediction using classifier models and SHAP analysis to identify important features. SHAP identifies features important for prediction of each fate label. SHAP correlation identifies whether the value of a feature is positively or negatively correlated with the output probability of a fate. (**b**) Boxplots showing accuracy values obtained with the three model architectures for either the RNA (left panel) or the ATAC (right panel) model. Overall, Random forest was the best performing architecture (best in ATAC and second best in RNA) and was chosen for all downstream analysis. (**c**) Same plot as Fig. 2j but for F1-weighted scores. (**d**) Left Panel: Boxplots showing variation in F1-weighted score values for ATAC models trained on full peak sets for 'all', 'distal', 'intronic', 'exonic' or 'promoter' peaks (Mann Whitney Wilcoxon test; p-values: **** = $p < 0.0001$). Right Panel: Same plot but only for 'distal', 'intronic', 'exonic' and 'promoter' peak sets, subsampled to the same number of peaks (n = 8823; Mann Whitney Wilcoxon test; p-values: **** = $p < 0.0001$; ** = $p < 0.01$; * = $p < 0.05$). (**e**) Heatmaps depicting mean TF activity scores for fate predictive TFs (as identified from SHAP analysis) across groups of state sisters. TFs show strong fate biased enrichment patterns in 'distal', 'intronic' and 'all' peaks but not exonic and promoter datasets. (**f**) Heatmap depicting Rank correlation of SHAP values for top predictive TFs shows high similarity between 'distal', 'intronic' and 'all' peaks models. (**g**) Bar plots of mean absolute SHAP values for a few TFs for fates as indicated. Bars are colored based on magnitude of SHAP correlation. SHAP analysis reveals that motif activity of many lineage specifying TFs is less predictive of cell fate in 'promoter' and 'exonic' models, while remains comparable across models for some others. Positive SHAP correlation for a feature in a given fate implies that higher values of the feature lead to higher probability of the model outputting that fate label. Negative correlation indicates lower values of the feature lead to higher probability of the model outputting that fate label.

#### Extended Figure 6

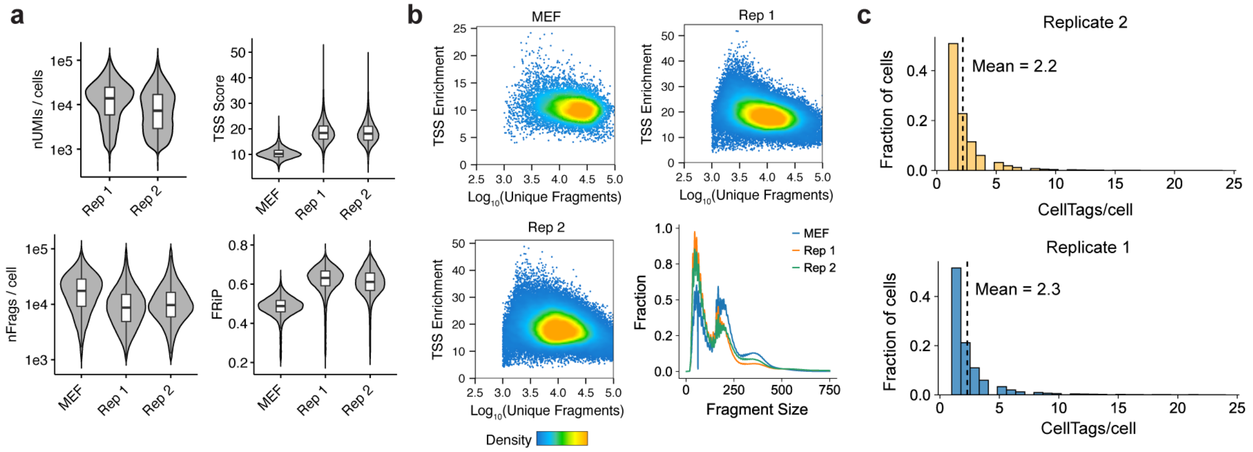


**d**

All clones

| | | Replicate 1 | Replicate 2 |
|---|---|---|---|
| Number of clones | | 3,068 | 5,416 |
| Number of RNA cells in clones | Day 3 | 4,781 | 11,537 |
| | Day 12 | 1,784 | 1,792 |
| | Day 21 | 3,492 | 8,923 |
| Number of ATAC cells in clones | Day 3 | 379 | 871 |
| | Day 12 | 676 | 675 |
| | Day 21 | 3,539 | 7,989 |

State-fate clones

| | | Replicate 1 | Replicate 2 |
|---|---|---|---|
| Number of clones | | 738 | 690 |
| Number of RNA cells in clones | Day 3 | 1,329 | 1,817 |
| | Day 12 | 1,223 | 1,126 |
| | Day 21 | 2,290 | 2,112 |
| Number of ATAC cells in clones | Day 3 | 131 | 266 |
| | Day 12 | 429 | 300 |
| | Day 21 | 1,064 | 373 |

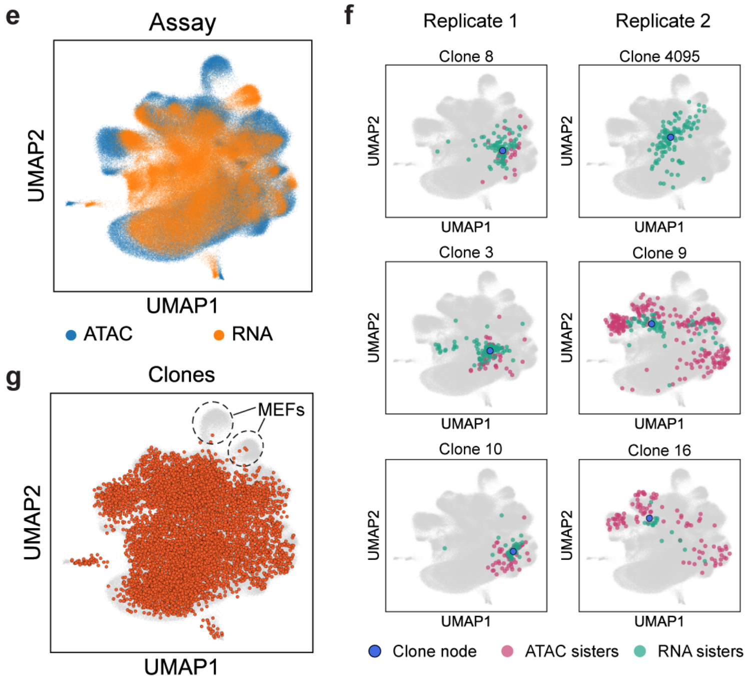


**Extended Figure 6. Single-cell metrics for the direct reprogramming dataset.** (**a**) Violin plots for single-cell quality metrics for the scRNA-seq and scATAC-seq datasets, split by biological replicates. (**b**) Unique fragments/cell vs single-cell TSS enrichment scatterplots and fragment size distribution plots for the scATAC-seq dataset. (**c**) Histograms of number of CellTags detected per cell across the two biological replicates after filtering and processing of CellTag reads. (**d**) Summary of all clones identified across single-cell modalities, for both biological replicates. (**e**) Cells in the clone-cell embedding UMAP with assay information projected shows uniform embedding of both single-cell modalities. (**f**) UMAPs depicting representative clone nodes from both biological replicates along with their constituent cells. (**g**) UMAP with all clone nodes highlighted shows uniform distribution of clones across all cell states except the unlabeled MEFs.

### Extended Figure 7

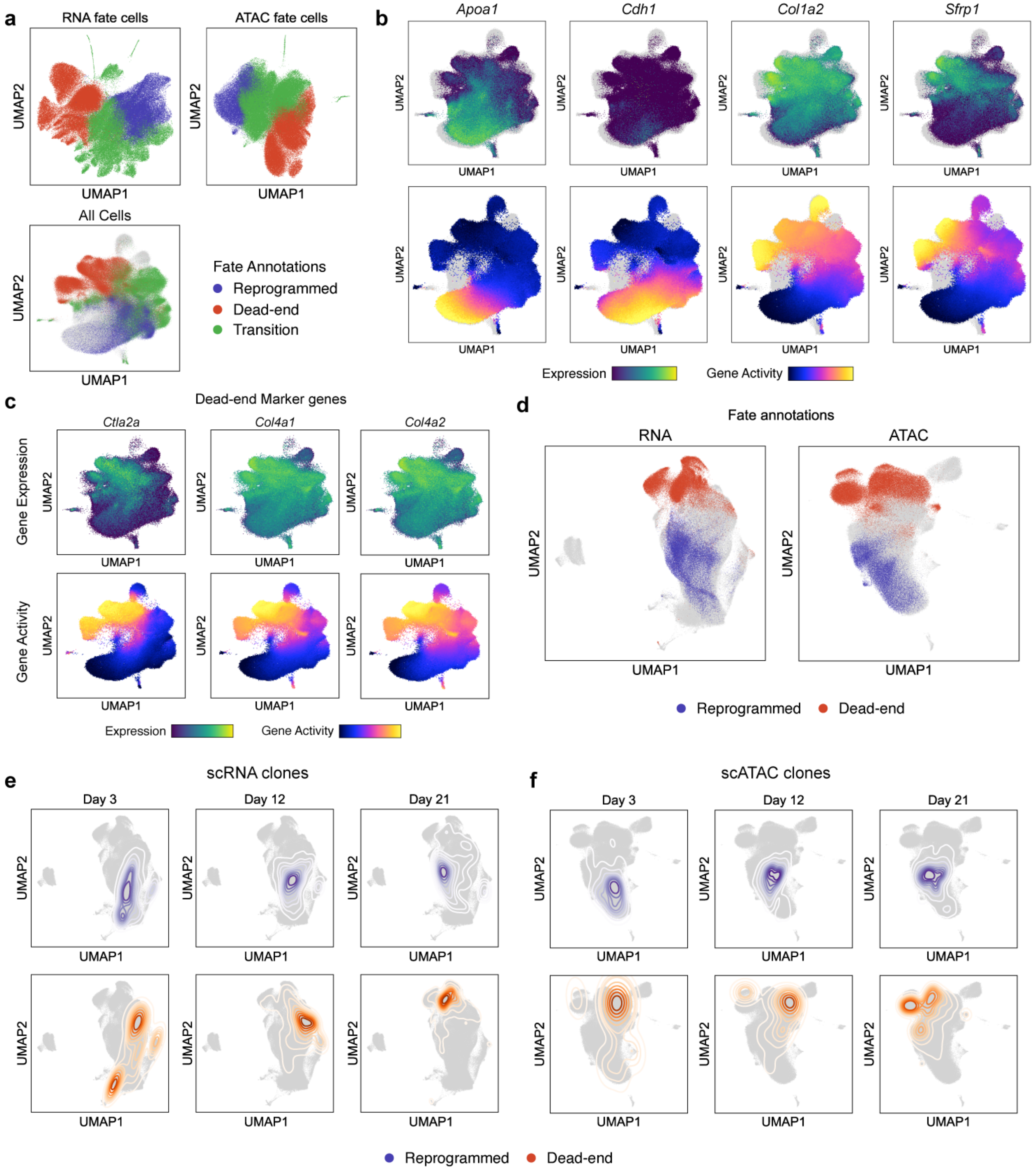

**Extended Figure 7. Fate annotation in direct reprogramming.** (**a**) UMAPs with ‘reprogrammed’, ‘dead-end’ and ‘transition’ fate information projected. Fate cells (Days 12 and 21) were re-clustered and annotated with one of the three fates based on marker gene expression/accessibility, in both modalities independently. (**b**) Clone-cell embedding UMAPs with expression and accessibility information for key marker genes projected. (**c**) UMAPs with expression of key dead-end marker genes projected. (**d**) UMAPs for individual modalities with reprogrammed and dead-end fate information projected. (**e**) Contour plots showing longitudinal tracking of cell fates enabled by CellTagging, independently for both scRNA and scATAC.

#### Extended Figure 8

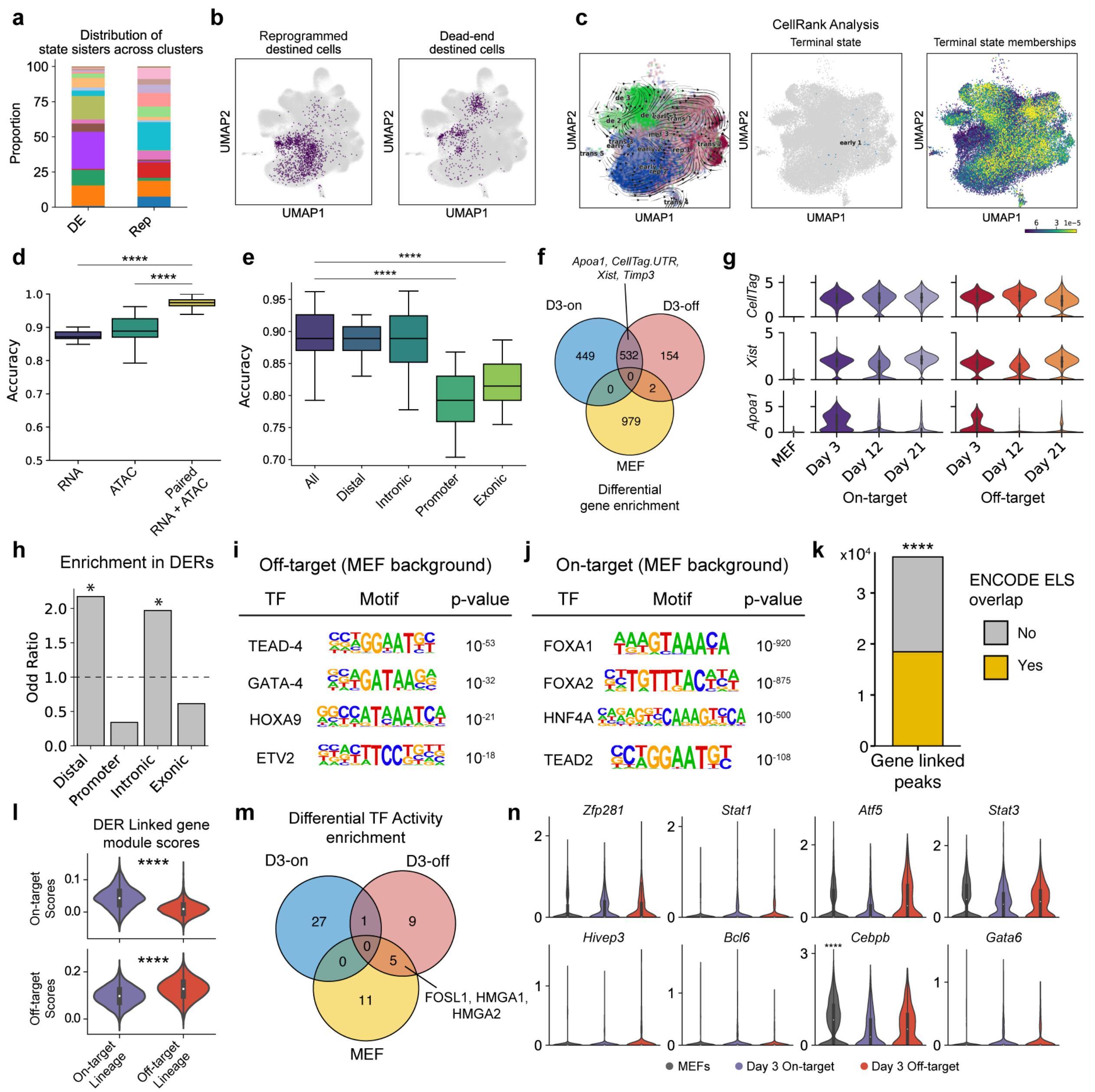


**Extended Figure 8. Differential analysis of expression and chromatin accessibility state across lineages.** (**a**) Distribution of reprogramming and dead-end destined cells across clusters and (**b**) their projection on the clone-cell embedding UMAP. (**c**) CellRank analysis fails to reveal true lineage dynamics underlying direct reprogramming. This analysis was run on a random subset of 40,000 scRNA-seq cells. Left Panel: Velocity vectors and RNA clusters overlaid onto the UMAP. Middle Panel: ‘Early_1’, a cluster from Day 3 cells identified as a terminal state by

CellRank. Right Panel: Continuous membership values for the terminal state 'Early_1'. (**d**) Fate prediction from Day 3 cell state using random forest classifiers. As expected, a combination of both gene expression and imputed TF activity scores outperforms either feature space individually (Mann Whitney Wilcoxon test; p-values: Paired vs ATAC = 3.5e-09; Paired vs RNA = 1.4e-09). (**e**) State-fate prediction analysis using functional subsets of accessible peaks reveals higher lineage priming in distal and intronic regions (Mann Whitney Wilcoxon test; p-values: All Peaks vs Promoter = 1.757e-08; All Peaks vs Exonic = 1.052e-07). (**f**) Venn diagram summarizing differentially enriched genes across uninduced MEFs and the two reprogramming fates on Day 3. This analysis identified 2,116 genes with varying degrees of overlap between groups (D3-on: Day 3 on-target cells; D3-off: Day 3 off-target cells). (**g**) Violin plots for several genes enriched in both reprogramming fates on Day 3. (**h**) DERs are enriched in distal and intergenic regions of the genome. (Fischer's exact test; p-values: 0 for both intronic and distal peaks). HOMER analysis to identify motifs enriched in (**i**) Off-target (dead-end) DERs and (**j**) On-target (reprogrammed) DERs, compared to a MEF DER background. (**k**) Enrichment of ENCODE cCRE Enhancer Like Elements in gene linked peaks. (Permutation test; 10,000 permutations, p-value: 1e-04). (**l**) Enrichment of DER linked genes' module scores in each lineage (Mann Whitney Wilcoxon test; p-values: top panel = 6.9e-306; bottom panel = 0). (**m**) Venn Diagram summarizing differential enrichment of 53 TF activities across uninduced MEFs and the two reprogramming fates on Day 3. (**n**) Violin plots showing expression of off-target associated TFs, as identified from TF activity analysis, across uninduced MEFs and the two reprogramming fates on Day 3. *Cdx1* expression was not detected in any of the groups and is hence not plotted. All TFs, except *Cebpb* (MEF enriched; adjusted p-value: 9.17e-16), did not show any lineage specific enrichment of expression.

#### Extended Figure 9

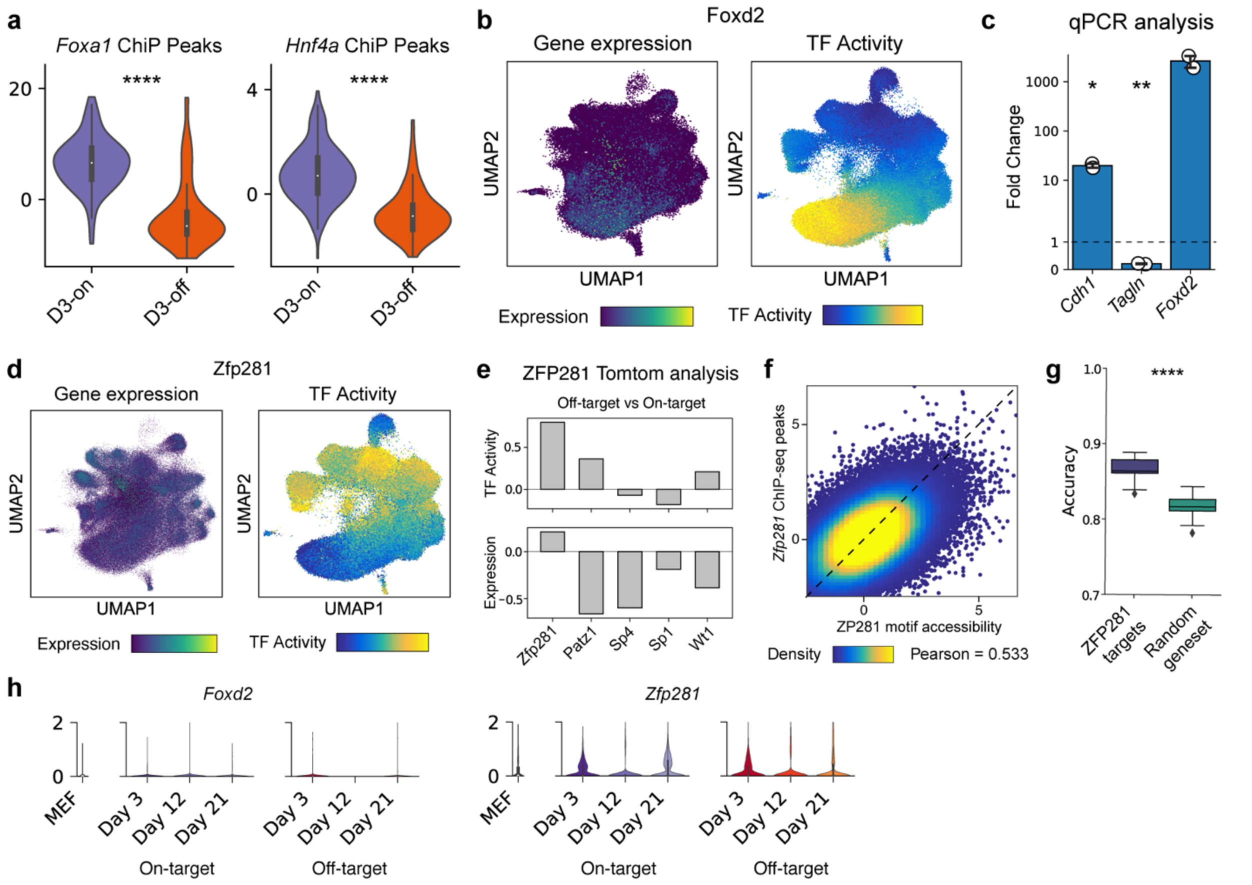


**Extended Figure 9. Identification of Zfp281 and Foxd2 as regulators of iEP reprogramming.** (**a**) Violin plots comparing accessibility z-scores of FOXA1 and HNF4A genomic binding sites across the two reprogramming fates on Day 3 (Mann Whitney Wilcoxon test; p-value: FOXA1 = 1.6e-22, HNF4A = 1.1e-2) suggesting higher on-target binding of the two TFs in the on-target reprogramming lineage on Day 3. (**b**) Projection of *Foxd2* gene expression and FOXD2 TF activity levels on the clone-cell embedding. (**c**) Bar plots showing fold-change in on-target and off-target marker genes (*Cdh1* and *Tagln* respectively) upon *Foxd2* over-expression, compared to a GFP control, on reprogramming day 12 (t-test; p-values: *Tagln* = 0.006, *Cdh1* = 0.03; n=2 biological replicates). (d) Projection of *Zfp281* gene expression and ZFP281 TF activity levels on the clone-cell embedding. (**e**) Tomtom analysis identified four dead-end enriched TFs with significantly similar motifs to ZFP281. ZFP281 shows the highest enrichment in dead-end cells for both gene expression and TF activity levels across all TF candidates. (**f**) Scatterplot showing correlation between single-cell accessibility of ZFP281 genomic binding sites and ZFP281 motifs (Pearson correlation coefficient = 0.533). (**g**) Boxplot showing significantly higher cell fate prediction accuracy using ZFP281 target genes (1,612 genes) compared to a size matched set of random genes (Mann Whitney Wilcoxon test; p-value = 2.248e-09). (**h**) Violin plots showing expression levels of *Foxd2* and *Zfp281* in uninduced MEFs and along the two lineages.

#### Extended Figure 10

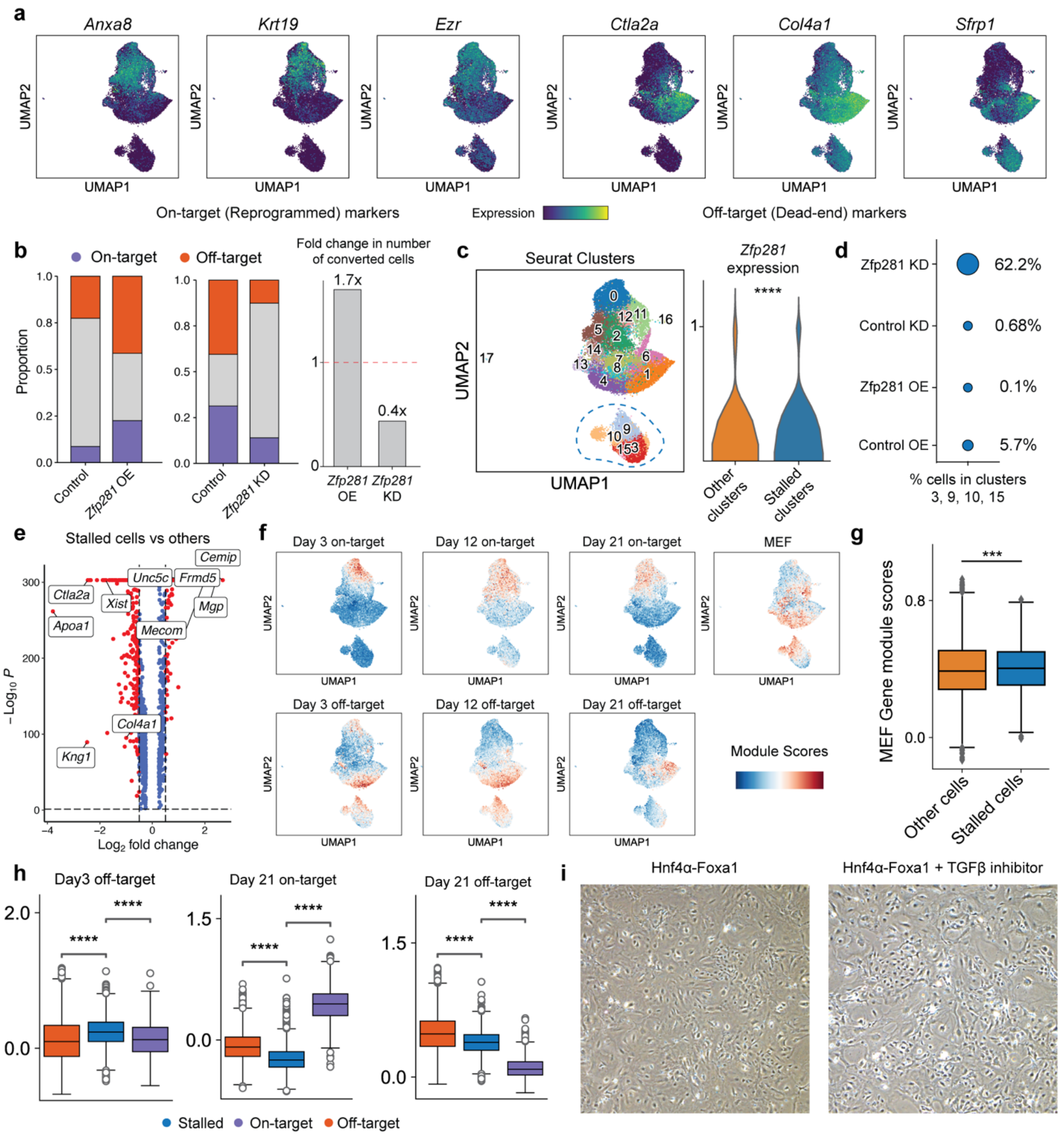


**Extended Figure 10. Single-cell analysis of *Zfp281* knockdown and overexpression.** (**a**) Projection of key on-target and off-target reprogramming marker genes on the UMAP for *Zfp281* overexpression and knockdown cells. (**b**) (Left Panel) Bar plots showing proportion of on-target and off-target fate cells and (Right Panel) change in total number of reprogrammed cells across the KD and OE experiments. A positive correlation between rate of reprogramming and *Zfp281*

expression suggests a role for the TF in promoting fate conversion away from the starting MEF identity. (**c**) (Left Panel) UMAP highlighting a distinct sub-population of cells, likely representing a stalled reprogramming cell state. (Right Panel) Violin plots comparing expression of *Zfp281* in the stalled cells versus rest of the cell population show that *Zfp281* expression is significantly lower in the stalled cells (Mann Whitney Wilcoxon test; p-value = 1.43e-15). (**d**) Dot plot showing the proportion of each sample in the stalled clusters. Cells from the *Zfp281* KD sample are enriched in the stalled cell states. (**e**) Volcano plot showing genes differentially enriched in the stalled cell sub-population (adjusted p-value < 0.05; Benjamini-Hochberg correction, absolute log2 fold-change > 0.5). (**f**) Gene expression module scores for MEF, on-target and off-target marker genes from all three time points, projected on the UMAP. (**g**) Box plot showing slight enrichment of MEF marker genes module scores in the stalled clusters (Mann Whitney Wilcoxon test; p-value = 0.00013). (**h**) Box plots comparing module scores for Day 3 off-target, Day 21 off-target, and Day 21 on-target marker genes module scores across stalled cells and the two reprogrammed clusters (Mann Whitney Wilcoxon test; **** = p-value < 0.0001). Marker genes for module scoring were obtained from our lineage analysis. (**i**) Brightfield microscopy images for reprogramming cells on Day 3 of reprogramming in presence (Right Panel) or absence (Left Panel; Vehicle) of TGF-β signaling inhibitor SB431542.
